# NOHIC: A PIPELINE FOR PLANT CONTIG SCAFFOLDING USING PERSONALIZED REFERENCES FROM PANGENOME GRAPHS

**DOI:** 10.64898/2026.03.17.712436

**Authors:** An Nguyen-Hoang, Kübra Arslan, Venkataramana Kopalli, Steffen Windpassinger, Dragan Perovic, Andreas Stahl, Agnieszka Golicz

## Abstract

Hi-C data is commonly used for reference-free *de novo* scaffolding. However, with the rapid increase in high-quality reference genomes, reference-guided workflows are now more practical for assembling large numbers of target genomes without relying on costly and labor-intensive Hi-C sequencing. Recently, a pangenome graph-based haplotype sampling algorithm was introduced to generate personalized graphs for target genomes. Such graphs have strong potential as references for reference-guided contig scaffolding. Here, we present noHiC, a reference-guided scaffolding pipeline supporting key steps of plant contig scaffolding. A distinctive feature of noHiC is the *nohic-refpick* script, generating a best-fit synthetic reference (synref) from a pangenome graph that is genetically close to the target contigs. This enables the integration of genetic information from many references (up to 48 in our tests) without using them separately during scaffolding. Synrefs showed advantages over highly contiguous conventional references in reducing false contig breaking during reference-based correction. Additionally, *nohic-refpick* can be combined with fast scaffolders (ntJoin) to rapidly produce highly contiguous assemblies using synrefs derived from pangenome graphs. The noHiC pipeline, used alone or in combination with ntJoin, can generally produce assemblies that are structurally consistent with public Hi-C-based or manually curated genomes. The pipeline is publicly available at https://github.com/andyngh/noHiC.

**GRAPHICAL ABSTRACT:** 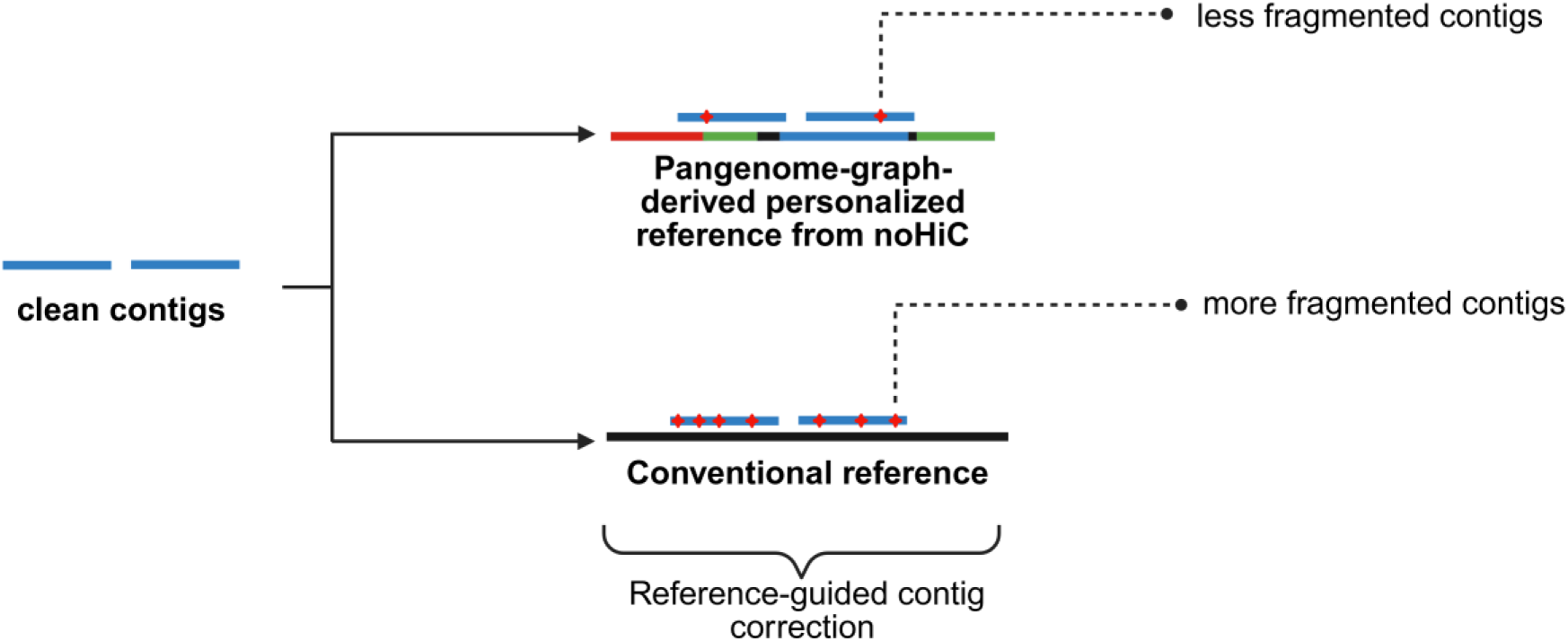

## INTRODUCTION

Despite challenges arising from the characteristics of plant genomes, including wide variation in genome size, ploidy level, heterozygosity, and high repeat content, the quality of plant genome assemblies has improved substantially in recent years (1, 2). Genome assembly workflows that integrate PacBio HiFi, Oxford Nanopore ultra-long (ONT UL), and Hi-C sequencing data are now capable of producing telomere-to-telomere (T2T) or near-T2T assemblies (3). These advances have provided plant genomics with high-quality genomes, including gapless T2T assemblies, enabling the construction of pangenome graphs that incorporate genetic information from numerous genome assemblies and thereby improving the understanding of genetic architectures (4). For example, complete T2T genomes for sorghum, lettuce, *Medicago*, and *Lablab purpureus* have recently been assembled using combinations of PacBio HiFi, ONT UL, and Hi-C sequencing technologies (5–8).

However, Hi-C-based reference-free scaffolding requires high sequencing depth, which is both costly and computationally intensive (9–12). In addition, Hi-C library preparation involves complex and time-consuming protocols that might require specific optimizations for a species of interest (10, 13, 14). In contrast, reference-guided genome scaffolding workflows, which use alignments between target and reference genomes to order and orient contigs (15), offer an effective means to reduce sequencing costs (12) and thereby support projects involving the assembly of tens to hundreds of target genomes. This approach has become increasingly practical given the rapid growth in both the number and quality of published plant genomes (1).

Reference-guided genome scaffolding can be subject to reference bias, where results are distorted at atypical (for example, highly divergent) positions on a reference genome (16). In particular, reference bias can negatively impact sequence alignment, as reads or sequences from diverse regions may be poorly mapped or fail to be mapped to a single reference genome (16, 17). Because most reference-guided assembly tools require alignment of target contigs to a reference genome (12, 18–20), divergent regions in the target genome may fail to be reconstructed during reference-guided assembly and may be absent from the final assembly, resulting in reduced representation of genetic diversity (15, 21–23). In addition, errors present in reference genomes can propagate to the target assemblies and result in misassemblies (15, 24). To address these issues, several tools have been developed for multi-reference scaffolding, including Ragout2, ntJoin, and Multi-CSAR (20, 25, 26). However, the algorithms implemented in these tools do not yet support efficient scaffolding with a high number of references (e.g., >10) or their repeated use. In particular, Ragout2 (for large genomes) requires a HAL/MAF (Hierarchical Alignment/Multiple Alignment Format) file produced by Cactus (27), which contains alignments between the target genome and the references. Consequently, users must undergo the complex process of updating the HAL file’s tree structure whenever new assemblies become available (https://github.com/ComparativeGenomicsToolkit/cactus/blob/master/doc/updating-alignments.md). Multi-CSAR constructs multiple separate single-reference draft assemblies for the target genome before generating a single multi-reference assembly (25), which is not time-efficient when many references are used. ntJoin relies on user-specified weights for each reference (26) (https://github.com/bcgsc/ntJoin), which must be re-optimized whenever a new assembly is required.

A pangenome represents the entirety of DNA sequences in a species, including both coding and non-coding regions (17, 28). Comparisons of individual genomes from the same species typically reveal a large number of variants, ranging from single-nucleotide polymorphisms (SNPs) to large structural variants (SVs) that can result in the absence of entire genes in an individual (28, 29). Thus, pangenomes provide an effective framework for compiling genetic variation from multiple genomes to address reference bias.

The haplotype sampling algorithm for pangenome graphs proposed by Sirén *et al.* has the potential to improve multi-reference-guided scaffolding. Given a pangenome graph, the algorithm can recombine haplotypes to create a personalized graph for a target genome by using graph-unique k-mers (i.e., k-mers occurring once in the graph) present in the target’s sequencing reads (30). Consequently, recombined DNA sequences originating from different haplotypes can be extracted and used as a single reference for scaffolding, without directly relying on multiple references. The limitations of current multi-reference scaffolders can therefore be addressed, as pangenome graphs can be reused to generate a single synthetic reference for new target assemblies without requiring multiple whole-genome alignments or reference weight optimization. However, no attempt has been made to apply pangenome graph-based haplotype sampling to reference-guided contig scaffolding to date.

Pangenome graphs have been generated for over 20 major crop species or genera, including cereals, legumes, vegetables, fruit, and tuber-bearing crops (17). Assemblies incorporated into plant pangenomes have been used primarily for variant-trait association studies, building repositories of resistance genes, and investigating wild relatives of crops (28). The application of the haplotype sampling algorithm in plant genome scaffolding may represent the next use of pangenome graphs, whereby pangenomes from publications or collections of public assemblies can be used to generate best-fit references for diverse target genome assemblies.

Here, we introduce the noHiC pipeline, which encompasses key steps in reference-guided scaffolding, including removal of contaminant contigs, correction of contig misassemblies, contig scaffolding using a pangenome graph-based or a conventional reference genome, and assessment of assembly quality through metrics and visualizations. Notably, noHiC incorporates the pangenome haplotype sampling algorithm in the sub-script called *nohic-refpick* to generate a personalized reference (synref) that is genetically close to the target genome and improves scaffold contiguity compared with those constructed from a single conventional reference. Across all tests, assemblies scaffolded using noHiC generally showed strong structural consensus with Hi-C-based or manually curated public assemblies. This advantage of noHiC was also evident when an alternative tool was used to scaffold target genomes with synrefs generated by the noHiC sub-script. Based on the results of our assembly evaluations, noHiC represents a promising pipeline for reference-guided contig scaffolding when pangenome graphs are accessible and/or Hi-C sequencing data are not available.

## MATERIAL AND METHODS

### noHiC Pipeline Overview

The noHiC pipeline includes four sub-scripts that can be executed independently, including *nohic-clean, nohic-refpick, nohic-asm*, and *nohic-eval*. HiFi reads, or other error-corrected long reads, are required for *nohic-refpick, nohic-asm*, and *nohic-eval*. Detailed instructions for pipeline execution, sub-scripts, and the versions of required dependencies are available at: https://github.com/andyngh/noHiC.

### Contaminant Contig Removal – nohic-clean

The removal of contaminant contigs is performed by the *nohic-clean* subscript, using raw contigs generated by tools such as hifiasm (31) and Canu (32) as inputs. *nohic-clean* consists of four steps and can be restarted from a previously failed step using the “--resume” flag.

The subscript begins by screening contigs for the presence of sequencing adapters, based on user-specified adapter sequences. If adapters are detected in the input data, *nohic-clean* halts and outputs the names and sequences of the contigs containing adapters. Users may remove these adapter-containing contigs before resuming *nohic-clean*, or they may restart contig assembly after performing more effective adapter trimming of the reads. After confirming that the contigs are free of adapters, *nohic-clean* calls Kraken2 (33) and Taxonkit (34) to assign taxonomic origins to the contigs. At this stage, users must provide the path to a Kraken2 database and specify the taxonomic group from which the target contigs originate (e.g., Viridiplantae) to distinguish contaminants from target sequences. In the subsequent optional step, users may choose to exclude contigs derived from organelles (mitochondria and chloroplasts) by supplying *nohic-clean* with reference organellar sequences. *nohic-clean* then uses BLASTn (35) to remove contigs showing ≥ 90% identity and query coverage relative to the reference organellar sequences.

### Personalized Reference Generation – nohic-refpick

To build a personalized reference (synref) for scaffolding a target genome, users must provide *nohic-refpick* with a FASTQ file containing error-corrected long reads, a .gbz file (36), and a .hapl file (30). The pangenome graph in GBZ format and its haplotype information (.hapl files) can be from previous publications or generated beforehand using pangenome graph builders such as Minigraph-Cactus (37).

Once the input data are specified, *nohic-refpick* first runs KMC (38) to count k-mers (default: 29 bp) in the reads and stores them in a KFF file (39). This file, together with the .gbz and .hapl files, is then supplied to *vg haplotypes* (30) to generate a personalized graph for the target genome using the default haplotype sampling algorithm. Briefly, the input pangenome graph is partitioned into 10-kb blocks, from which graph-unique k-mers, occurring only once in the graph and effectively representing the incorporated haplotypes, are extracted. The algorithm then queries these graph-unique k-mers against KFF files generated from reads of the target genome. Based on k-mer counts and coverage, which are automatically estimated by *vg haplotypes*, the graph-unique k-mers detected in the KFF file are classified as homozygous, heterozygous, or absent. In the final step, the best-fitting haplotypes from each 10-kb block are selected according to the k-mer classification profile, resulting in a personalized graph (30). Extraction of the synref from the personalized graph is performed using *vg paths* (--extract-fasta). The resulting synref is an artificial recombinant reference genome generated by combining the best-fitting 10-kb blocks from different haplotypes within the input pangenome graph. Users may either use the synref directly at this stage or further refine it (by gap patching) before employing it as a reference for reference-guided scaffolding of the target genome.

To improve the quality of the synref, users may optionally enable gap patching with the --patch (-p) flag, which starts the process of replacing gaps in the synref with sequences from a donor genome. Ideally, the donor genome used for synref patching is a gapless, T2T assembly. However, users may select any genome with the highest available contiguity within their species of interest as the donor. In patching mode, an in-house script (“*Asm_Decomposing.sh*”) is executed to decompose the synref into contigs at gap positions. These synref contigs are then aligned to a user-specified high-quality donor genome using minimap2 (40). The resulting SAM file is converted to BAM format with *samtools view* (41), and the BAM file is subsequently used as input for GPatch (42) to fill the gaps using sequences from the donor genome.

### Contig Error Correction and Scaffolding – nohic-asm

The correction of contig misassemblies and subsequent scaffolding is performed by *nohic-asm*, which consists of five computational steps: chimeric contig breaking based on clipped reads (i.e., reads that are only partially mapped), correction of small misassemblies, coverage-based chimeric contig breaking, scaffolding, and gap closing. All steps are optional except scaffolding. As with *nohic-clean*, *nohic-asm* can be resumed from a previously failed step.

*nohic-asm* calls CRAQ (43) in the first step of the pipeline to break chimeric contigs by mapping input long reads to the input contigs and identifying clipped reads as break signals. The default minimum proportion of clipped reads required to break a contig is 75%. Users may optionally choose to break heterozygous misjoins by applying the --ignore-het flag, which sets CRAQ’s --sms_clip_coverRate parameter to 55%.

In the second step, Inspector (44, 45) is used to correct small misassemblies in the contigs (base substitutions, expansions, collapses, and inversions) using the input long reads. Remaining chimeric contigs are then broken using *ragtag.py correct* (18). Putative breakpoints are identified by aligning the input contigs to a reference genome (either the synref or another reference) and validated by detecting abnormal increases or decreases in read coverage inside a window (which can be 10 or 45 kb long). The upper and lower coverage thresholds used to validate breakpoints are automatically determined by RagTag. *nohic-asm* provides five correction presets for this step, with the following parameters for *ragtag.py correct*:

**draft**: --aligner minimap2 --mm2-params "-x asm5" -v 10,000

**luck**: --aligner minimap2 --mm2-params "-x asm5" -v 45,000 --remove-small

**standard** (default): --aligner nucmer --nucmer-params "--maxmatch -l 100 -c 500" -v 45,000 --remove-small

**aggressive**: “standard” parameters plus -d 50,000

**raw**: “standard” parameters without read validation and --remove-small

Because minimap2 allows alignments between noisy query and target (reference) sequences through short exact matches (19-bp minimizer seeds) under the “seed-chain-align” paradigm (40, 46), its use in *ragtag.py correct* results in lower misassembly detection granularity compared with Nucmer (using -l 100). Consequently, the “draft” and “luck” options in *nohic-asm* are considered relaxed correction presets, as they generate fewer contig breaks than the other presets.

The “draft” preset applies only the default parameters of *ragtag.py correct* and is intended for users who aim to perform an initial assessment of assembly quality. The “luck” preset is also a relaxed correction mode but employs a larger window size (45 kb) for read-based validation of misassemblies, thereby increasing sensitivity to misjoin detection. In addition, the --remove-small flag is enabled in the “luck” preset to exclude alignments shorter than 1 kb between the target and reference genomes. This reduces the likelihood that small alignments will incorrectly bridge larger syntenic regions (https://github.com/malonge/RagTag/wiki/correct). Under favorable conditions, the “luck” preset can correct most chimeric contigs while minimizing unnecessary breaks and preserving overall assembly contiguity, which motivated its designation.

Presets based on Nucmer (“standard”, “aggressive”, and “raw”) use longer exact seed matches (minimum length of 100 bp) for alignments between target and reference assemblies (47). As a result, alignments are divided into a greater number of fragments, increasing the number of candidate misassemblies subjected to read-based validation and, consequently, producing more contig breaks. These Nucmer-based presets enforce closer conformity between target and reference sequence arrangements and are intended for cases in which the reference genome is closely related to the target or when chimeric contigs cannot be resolved using relaxed presets. The “aggressive” preset applies a shorter maximum alignment merging distance (50 kb) than the “standard” preset (100 kb), leading to more unmerged alignments that are treated by *ragtag.py correct* as putative misassemblies (18). The “raw” preset uses alignment parameters similar to those of the “standard” preset but omits read-based validation, resulting in the breaking of all identified putative misassemblies.

The corrected contigs are then oriented into scaffolds using *ragtag.py scaffold* (18). Gaps are closed using TGSGapcloser with the --ne option (48).

### Assembly Quality Evaluation – nohic-eval

The *nohic-eval* sub-script enables users to assess scaffolded assemblies across several criteria, including assembly contiguity, gene content completeness, and structural correctness. *nohic-eval* consists of five evaluation steps. First, contiguity metrics (e.g., N50, auN, number of gaps) and scaffold lengths are calculated using *gfastats* and *bioawk* (49) (https://github.com/lh3/bioawk). BUSCO is then run with a user-specified lineage to evaluate the completeness of gene architecture (50). In the third and fourth steps, regional and structural assembly quality indices (R-AQI and S-AQI) and the quality value (QV) are computed using CRAQ (43) and Inspector (44) to assess structural correctness. AQI metrics are calculated as one hundred times the exponential of negative 0.1N divided by L, where N represents the cumulative normalized counts of CREs (clip-based regional errors) or CSEs (clip-based structural errors), and L denotes the assembly length in megabases. In the context of noHiC, which does not require short reads for scaffolding, CREs correspond to gaps or regions spanned by long reads containing mismatches, whereas CSEs refer to regions characterized by clipped long reads (43). QV is defined as minus ten times the base-ten logarithm of E, where E is equal to the total number of bases involved in assembly errors divided by the assembly length in base pairs (44). In the final step of *nohic-eval*, an in-house R script, *nohic-viz.R*, is used to generate an error map showing misassemblies identified by CRAQ and Inspector across chromosomes. Synteny between the target assembly and a reference genome can also be visually assessed by a dot plot using *paf2dotplot* (https://github.com/moold/paf2dotplot). Evaluation steps 2-5 are optional. As with *nohic-clean* and *nohic-asm*, users may resume *nohic-eval* from a previously failed step.

### Sequencing Data, Genome Assemblies, and Pangenome Collections

HiFi reads from four plant species were used to generate contigs for testing the noHiC pipeline: *Arabidopsis thaliana* CAMA-C-2 (SRA: ERR10084604) (51), *Sorghum virgatum* (SRA: SRR33597079) (52), *Glycine max* cv. Wm82-ISU-01 (SRA: SRR29929794 and SRR29929795) (53), and *Hordeum vulgare* cv. Foma (SRA: ERR10665167, ERR10665168, ERR10665169, ERR10665170, and ERR10665171) (54). The original public assemblies for these four species were also used as controls for comparison with assemblies produced by noHiC.

Four collections of 48, 11, 21, and 10 public assemblies were used to construct pangenome graphs for *A. thaliana* (*Ath*), *S. bicolor* (*Sbi*), *G. max* (*Gma*), and *H. vulgare* (*Hvu*), respectively, using Minigraph-Cactus (cactus v2.9.9) (37). The assembly of *S. virgatum* (*Svi*) was scaffolded using the *Sbi* pangenome and reference (51, 54–69). Within each assembly collection, one assembly was selected as a conventional (or ordinary) reference (i.e., a publicly available reference that is traditionally used as an input for reference-guided scaffolders) to guide contig scaffolding with noHiC. Accession numbers for all public assemblies are provided in **Supplementary Table S1**.

In the case of *Sbi*, the ordinary reference was not included in the pangenome graph collection, as this collection was also used in the assembly tests below involving our three in-house sorghum accessions to evaluate the performance of synrefs relative to a closely related conventional reference.

To assess the reusability of a pangenome graph in contig scaffolding, we used three of our *Sbi* accessions (named SB14122, B108, and ORE-18-14) and the mentioned pangenome graph containing the set of 11 public sorghum assemblies (55–58, 67) (**Supplementary Table S1**).

### Contig Assembly

Before contig assembly, we randomly sampled from the four plant species mentioned above a HiFi read subset of 5-7X coverage for each species and used these subsets only during the scaffold assembly evaluation process by *nohic-eval*. The read subsets were generated with seqkit v2.10.1 (70) using a fixed random seed (3108). These subsets were not used in any other sub-scripts of noHiC requiring long reads as inputs. This approach avoided bias in comparisons, as *nohic-asm* uses long reads for contig correction, and original public assemblies were used in comparisons. The evaluation subsets corresponded to 10% of reads from *Ath* CAMA-C-2, 10% of reads from *Svi* 1.0, all reads from sequencing run ERR10665168 for *Hvu* cv. Foma, and 25% of reads from *Gma* cv. Wm82-ISU-01 (SRA: SRR29929795). Contigs for each species were then assembled using the remaining HiFi reads as input and without Hi-C data. Hifiasm v0.25.0-r726 (or v0.16.1-r375 for *Gma*) was used to assemble the contigs (31).

In the evaluation of pangenome graph reusability for synref generation in the three *Sbi* genomes, no public assemblies were used as controls. Therefore, the full HiFi read sets were used to assemble contigs with Hifiasm v0.25.0-r726 (or v0.19.5-r590) (31).

### Pangenome Graph’s Reusability for Scaffolding

As described above, we constructed a small pangenome graph consisting of 11 *Sbi* assemblies and used it to generate synrefs for three *Sbi* target genomes (SB14122, B108, and ORE-18-14) to assess the reusability of pangenome graphs to build synrefs in contig scaffolding.

Contaminant and organellar contigs were removed using *nohic-clean* (parameters: -tg Viridiplantae -ioc yes -m no -kdb core_nt). PacBio adapter sequences used in this step are available at https://github.com/andyngh/noHiC. Kraken2’s *core_nt* database was used for taxonomic classification (https://benlangmead.github.io/aws-indexes/k2). Chloroplast and mitochondrial genome sequences from NCBI of *Sbi* (NC_008602.1, NC_008360.1) (71) were provided via the -ros argument.

Synref generation with *nohic-refpick* (-m 182 -k 29) was carried out with HiFi reads and pangenome graph indices (.gbz and .hapl) as inputs. For synref patching, the Hongyingzi genome (assembly ID: GCA_033546955.1) was used for the synref of SB14122 (55), while B108 and ORE-18-14’s synrefs were patched using the *Sbi* cv. CHBZ genome (assembly ID: GCA_040267525.1) (57).

To scaffold the clean contigs, *nohic-asm* was executed with two read-based correction presets, “luck” and “standard”, for each target genome to determine the conditions under which the advantages of synrefs were most apparent. Other parameters were as follows.

--run-gap-closing yes --craq-params "-x map-hifi" --inspector-params "--datatype hifi" --inspector-correct-params "--datatype pacbio-hifi" --ragtag-correct-params "-T corr" --ragtag-scf-params "-C -r -g 2" --tgsgapcloser-params "--tgstype pb"

The sequencing coverage values used for scaffolding were 20, 16, and 20 for SB14122, B108, and ORE-18-14, respectively. Three different references were used for each target genome: BTx623 v5 (58) (the conventional reference most closely related to the targets); the gapless donor reference used for synref patching; and the patched synref. This resulted in six assemblies per target genome (two presets × three references). After scaffolding, *nohic-eval* (-p hifi -b embryophyta_odb12) was used to assess the quality of the resulting assemblies.

Whole genome alignments between the three target genomes and BTx623 v5 were used to construct dot plots for misassembly visualizations. Putative misassemblies detected by the dot plots were confirmed by read mapping using IGV 2.16 (72). Contiguity metrics of the assemblies were calculated only for pseudo-chromosomes before gap closing. Metrics showing assembly correctness (AQIs and QV) were calculated for the whole assemblies after gap closing.

The computational test summary (Test 1) was illustrated in **Figure 1**.

**Figure 1:**
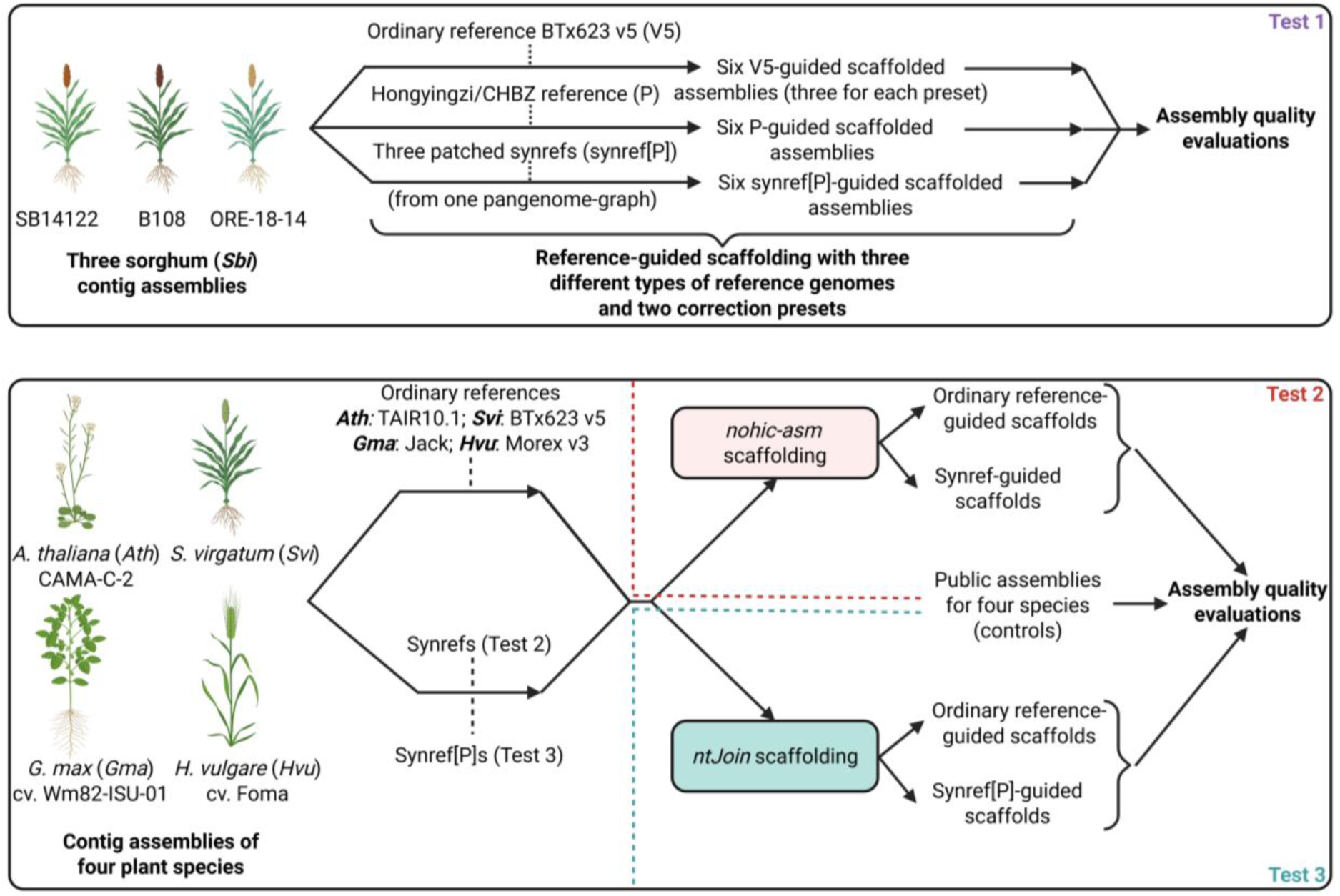
Summary of the three computational tests conducted in this study. Test 1 examines the reusability of a pangenome graph for generating multiple synrefs and identifies the correction preset in noHiC that best highlights the advantages of synrefs. Test 2 evaluates the effects of synrefs on contig scaffolding across four plant species. Test 3 assesses the flexibility of noHiC sub-scripts when combined with alternative scaffolding tools (ntJoin). The figure was created with BioRender (https://BioRender.com/70x3u2c).

### Comparing Assemblies Guided by Synref and Ordinary Reference in Different Species

To compare the quality of assemblies scaffolded using synrefs versus conventional references in different plant species, we applied the noHiC pipeline to contigs from four species (*Ath* CAMA-C-2, *Svi*, *Gma* cv. Wm82-ISU-01, and *Hvu* cv. Foma). The corresponding original public assemblies generated with Hi-C data or manual curation served as controls (51–54).

Contig decontamination was performed using *nohic-clean* as described in the previous computational tests. Chloroplast and mitochondrial genome sequences from NCBI for *Ath* (NC_037304.1, NC_000932.1) (69), *Sbi* (NC_008602.1, NC_008360.1) (71), *Gma* (CM021415.1, CM010429.1) (60), and *Hvu* (NC_008590.1, NC_056985.1, NC_042692.1, AP017300.1, AP017301.1) (71, 73, 74) were supplied to enable removal of contigs originating from organelles.

The sub-script *nohic-refpick* was used to generate unpatched synrefs for the four assemblies (-m 182 -p no -k 29) using HiFi reads and species-specific pangenome graph indices. The resulting clean contigs were then corrected and scaffolded for each species with two reference types using *nohic-asm*: a conventional reference genome (**Table 1**) and the corresponding unpatched synref. The *nohic-asm* script was executed using the “luck” correction preset. All other *nohic-asm* parameters were identical to those described above.

**Table 1.**
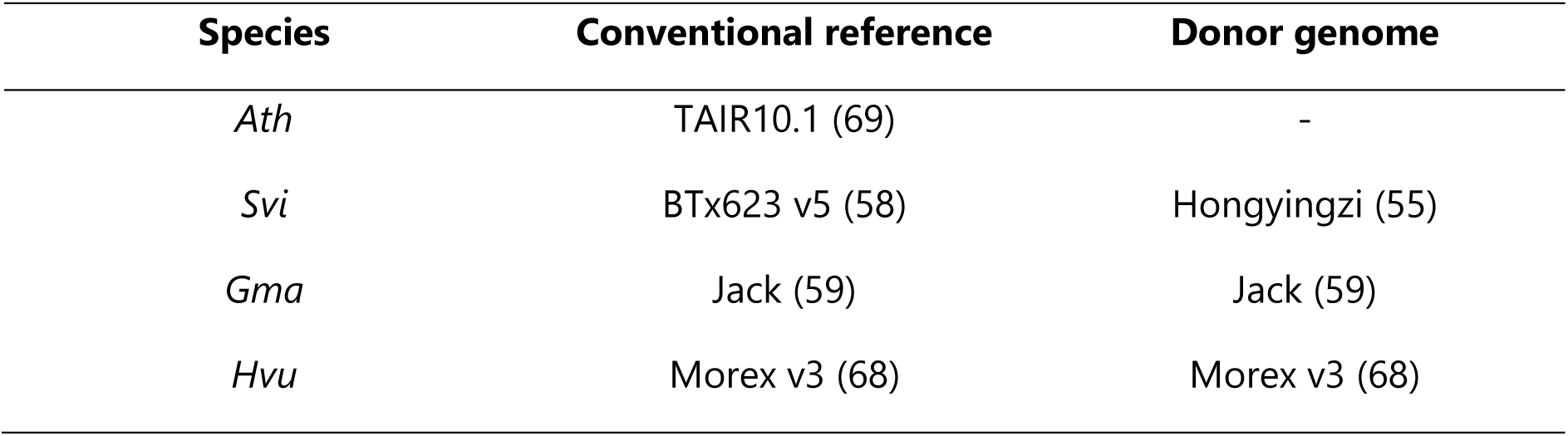
Conventional reference assemblies and donor genomes for synref patching of *Ath, Svi, Gma,* and, *Hvu*.

Sequencing coverage values provided to *nohic-asm* were 63, 44, 23, and 43 for *Ath*, *Svi*, *Hvu*, and *Gma*, respectively. Assembly quality was evaluated using the *nohic-eval* sub-script as described above. Whole genome alignments between each noHiC-based assembly and its control (original public assembly) were used for building dot plots to detect misassemblies. For each species, three assemblies were evaluated: the original public assembly and the two assemblies produced by noHiC.

The summary for this computational test involving the four plant species (Test 2) was illustrated in **Figure 1**.

### Combining noHiC Sub-scripts with a Fast Scaffolder

To demonstrate the flexibility of noHiC sub-scripts when used in combination with other scaffolders, clean contigs (outputs of *nohic-clean*) from *Ath* CAMA-C-2, *Svi*, *Gma* cv. Wm82-ISU-01, and *Hvu* cv. Foma were scaffolded using ntJoin with the following parameters: reference_weights=’2’ g=1 k=32 w=1000 n=2 (with w=500 for *Ath* CAMA-C-2) (26). The conventional references used for ntJoin are listed in **Table 1**.

Because ntJoin breaks contigs without read-based validation, *nohic-refpick* was executed with patching using donor genomes (**Table 1**) in cases where synrefs contained more gaps and/or exhibited lower contiguity than conventional references. This ensured that the potential benefits of synrefs were not underestimated due to false contig breaks. Generation of synrefs and assembly evaluation (for the whole assemblies) was performed by *nohic-eval* as described above.

The summary for this ntJoin-based computational test (Test 3) was illustrated in **Figure 1**.

### Genetic Distance between Synrefs and Target Genomes

In parallel with scaffolding, Mash v2.3 (75) was used with default parameters to calculate pairwise genetic distances among each clean contig assembly, its synref, its conventional reference, and the other assemblies within its respective pangenome collection. The resulting pairwise distance matrices were then used to construct Neighbor-Joining (NJ) trees with the *ape* and *ggtree* R packages (76, 77) to visualize the effectiveness of *nohic-refpick* in generating the reference most closely related to each target genome.

### Computational Resources

The noHiC pipeline was primarily developed and tested on computing nodes at the Bioinformatics Core Facility of Justus Liebig University Giessen within the German Network for Bioinformatics Infrastructure (de.NBI). These nodes operated under Linux kernels 6.1.0-23-amd64, 6.1.0-26-amd64, 6.1.0-31-amd64, 6.1.0-37-amd64, 6.1.0-38-amd64, 6.1.0-40-amd64, 6.12.32+bpo-amd64, or 6.12.33+deb12-amd64 and were managed using the Slurm job scheduling system. The available hardware resources ranged from 56 to 256 CPU cores and from 498 to 1992 GB of memory. In addition, a de.NBI virtual machine was used during pipeline development, running a 64-bit Ubuntu operating system with a Linux 6.8 kernel, 28 CPU cores, and 251 GB of memory.

### AI Use Disclosure

The core workflows of noHiC, which underpin the results presented in this study, were designed and structured using publicly available tools without the use of AI in computational tasks. The command-line interface (CLI), automation features, output structure, and several helper scripts of the noHiC pipeline (*nohic-viz.R* and *Asm_Decomposing.sh*) were conceptualized by the first author and implemented with assistance from ChatGPT5 (OpenAI). These components were manually tested to ensure the correct execution of the pipeline.

The manuscript was proofread with the assistance of ChatGPT5. The content and structure of the manuscript were prepared by the authors without AI involvement.

## RESULTS

### The noHiC Pipeline

The noHiC pipeline encompasses all essential steps in reference-guided contig scaffolding through four sub-scripts (or sub-pipelines). An overview of the main steps is presented in **Figure 2**.

**Figure 2:**
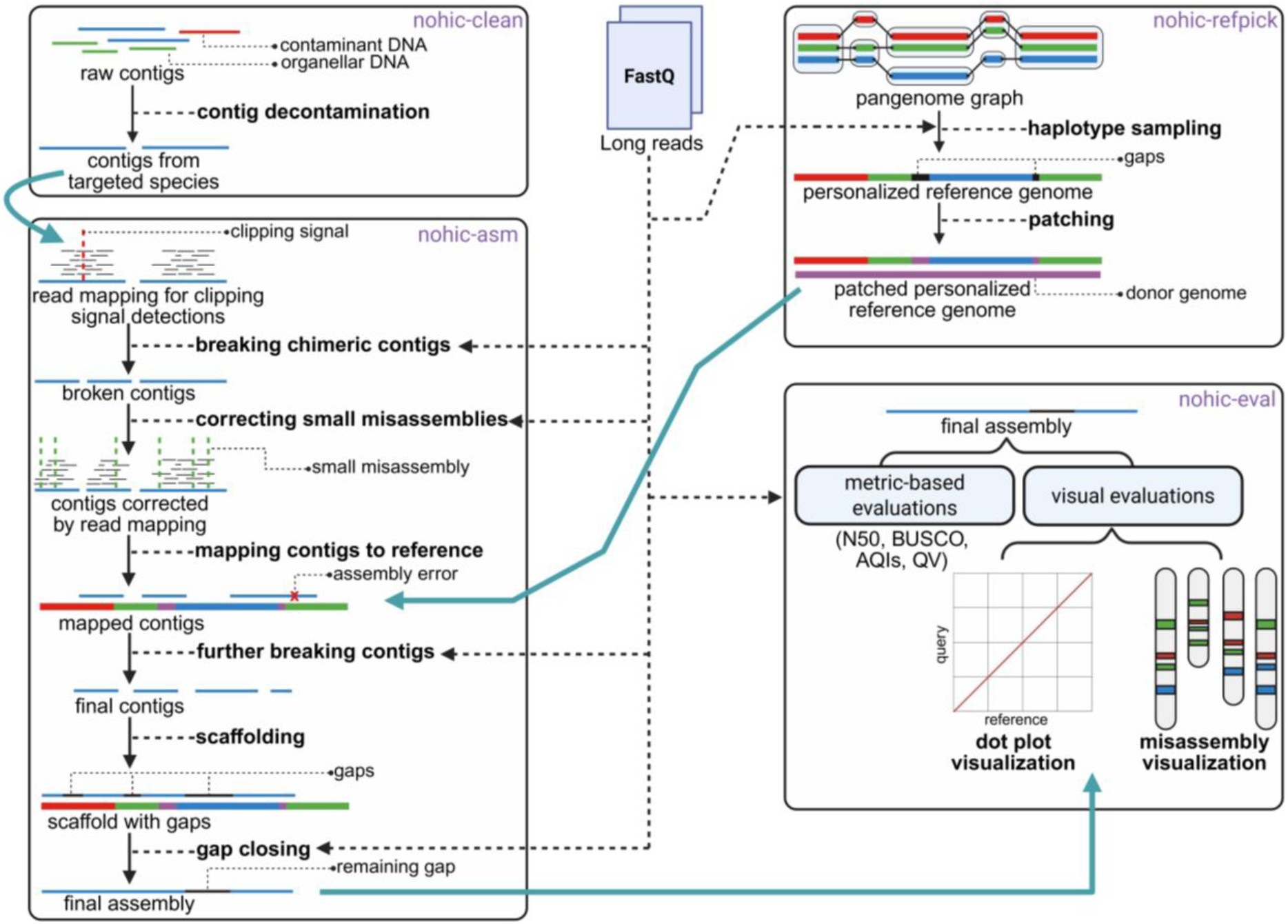
Diagram illustrating the computational tasks performed by the four sub-pipelines of noHiC. Cyan arrows indicate subsequent steps in which the final output of a sub-pipeline is used. Dashed arrows denote the steps in the pipeline that incorporate long reads. The figure was created with BioRender (https://BioRender.com/gpz1mxx).

The primary objective underlying the design of noHiC is to mitigate the adverse effects of reference bias in reference-guided contig scaffolding, thereby producing assemblies that are both contiguous and structurally accurate.

To accomplish this objective, the first step in the noHiC pipeline is to ensure that contaminant sequences present in the contig-level assembly do not compromise the structural integrity of subsequent scaffolding. Accordingly, *nohic-clean* is executed at the outset to perform taxonomic classification of contigs and remove contaminants (and, if desired, organellar DNA) prior to scaffolding.

To reduce reference bias, *nohic-refpick* was created to select the most appropriate haplotype sequences from a pangenome graph and combine them into a synthetic personalized reference genome (synref) for scaffolding the target contigs, using the haplotype sampling algorithm described by Sirén *et al.* (30). This approach ensures that the reference genome exhibits a high similarity in sequence organization to the target, reducing the likelihood of inaccurate contig correction and placement in downstream steps.

After preparing clean contigs and the synref, *nohic-asm* is used to polish the contigs and scaffold the corrected sequences into chromosomes. Three complementary strategies are implemented to ensure structural accuracy: clipped-read-based detection and breaking of chimeric contigs, correction of small misassemblies, and reference-guided correction to resolve remaining chimeric contigs. Corrected contigs are then oriented and concatenated into scaffolds according to the reference sequence (ideally, the synref, although a conventional reference may also be used), and gaps are subsequently closed.

Because a high-quality assembly must be both contiguous and accurate, *nohic-eval* provides a range of metrics and visualizations to assess assembly quality. Calculated contiguity metrics include N50; auN, the weighted sum of all Nx values (x from 0 to 100) (78, 79); number of gaps; and total contig and scaffold lengths. These metrics are used to evaluate contig fragmentation and chromosome anchoring. Gene space completeness is assessed using BUSCO. Structural accuracy is assessed through metrics such as the structural and regional assembly quality indices (S-AQI and R-AQI) and the quality value (QV). Visualizations illustrate misassembly locations and syntenic relationships between the target genome and a high-quality reference.

### Reusability of Pangenome Graphs for Generating Synrefs that Preserve Contig Contiguity

We collected 11 public *S. bicolor* (*Sbi*) assemblies (**Supplementary Table S1**) and constructed a pangenome graph, which was subsequently used to generate synrefs for scaffolding three in-house sorghum accessions, namely SB14122, B108, and ORE-18-14. The *nohic-asm* sub-pipeline was executed using two presets (“luck” and “standard”) and three reference types for each target assembly: the patched synref, BTx623 v5 (standard reference), and the donor genome used for synref gap patching (NCBI assembly IDs: GCA_033546955.1 – Hongyingzi or GCA_040267525.1 – CHBZ). For clarity, these reference genomes are referred to in this subsection and related ones as synref[P], V5, and P, respectively. V5 is genetically closer to the target genomes than the other assemblies used to construct the pangenome graph. In the case of ORE-18-14, one assembly in the pangenome collection, *S. bicolor* 654 T2T (GCA_048338455.1), is nearly as closely related to the target accession as V5. The two donor genomes in the P reference category are gapless assemblies.

The “luck” preset applies relatively relaxed contig correction, whereas the “standard” preset enforces stricter correction by constraining contig structure to more closely follow the reference. In total, six assemblies were generated for each *Sbi* accession (two presets × three references). This analysis was designed to address the following questions: (i) whether the benefits of synrefs in scaffolding are retained when a single pangenome graph is reused across multiple assemblies; (ii) whether a synref[P] performs better than the genetically closest assembly for target genome scaffolding; (iii) whether a synref[P] outperforms the gapless P references; and (iv) whether the advantages of synrefs are more evident under relaxed or strict correction presets.

The first observed positive effect of *nohic-refpick* was the generation of synrefs that closely match their target genomes. Specifically, the synrefs and synref[P]s for SB14122, B108, and ORE-18-14 are genetically closer to their respective targets than V5 and the other assemblies included in the pangenome collection, as shown by the Neighbor-Joining (NJ) trees in **Figure 3**. These results demonstrate that patching synrefs with donor genomes does not significantly alter the genetic relationship between a synref and its corresponding target assembly.

**Figure 3:**
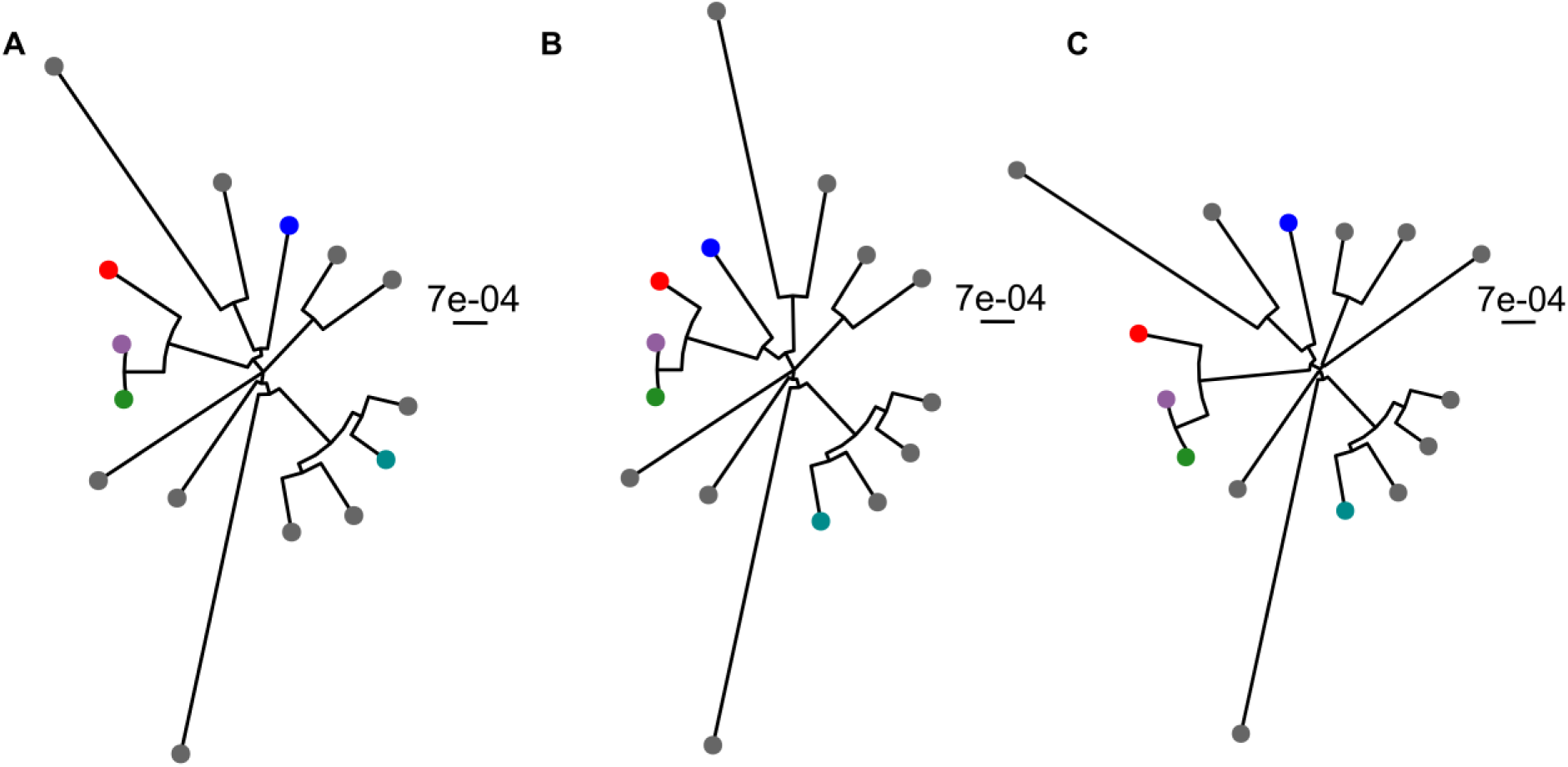
NJ tree constructions including the target *Sbi* assemblies SB14122 (A), B108 (B), and ORE-18-14 (C) (red dots), the corresponding synrefs (lavender dots), synref[P]s (green dots), V5 (blue dots), P references (cyan dots), and the remaining assemblies in the pangenome collection (gray dots). In all three cases, the synrefs and synref[P]s are the assemblies most closely related to the target.

In-chromosome contig contiguity metrics (i.e., the contiguity metrics for contigs placed in pseudo-chromosomes after scaffolding) prior to gap closing, including total contig length (Mb), contig number, contig auN (Mb), and gap number, were calculated for all six assemblies generated for each target *Sbi* accession.

To identify the correction preset that best highlights the benefits of synref[P]s, percentage changes in contig contiguity metrics were calculated between synref[P]-based assemblies and their corresponding V5- or P-based counterparts by subtracting the V5- or P-based values from the synref[P]-based values, dividing the change by the V5- or P-based values, and multiplying the result by 100. For each preset, this yielded six comparisons per metric, corresponding to three target assemblies and two reference contrasts (synref[P] versus V5 and synref[P] versus P) (**Figure 4**).

**Figure 4:**
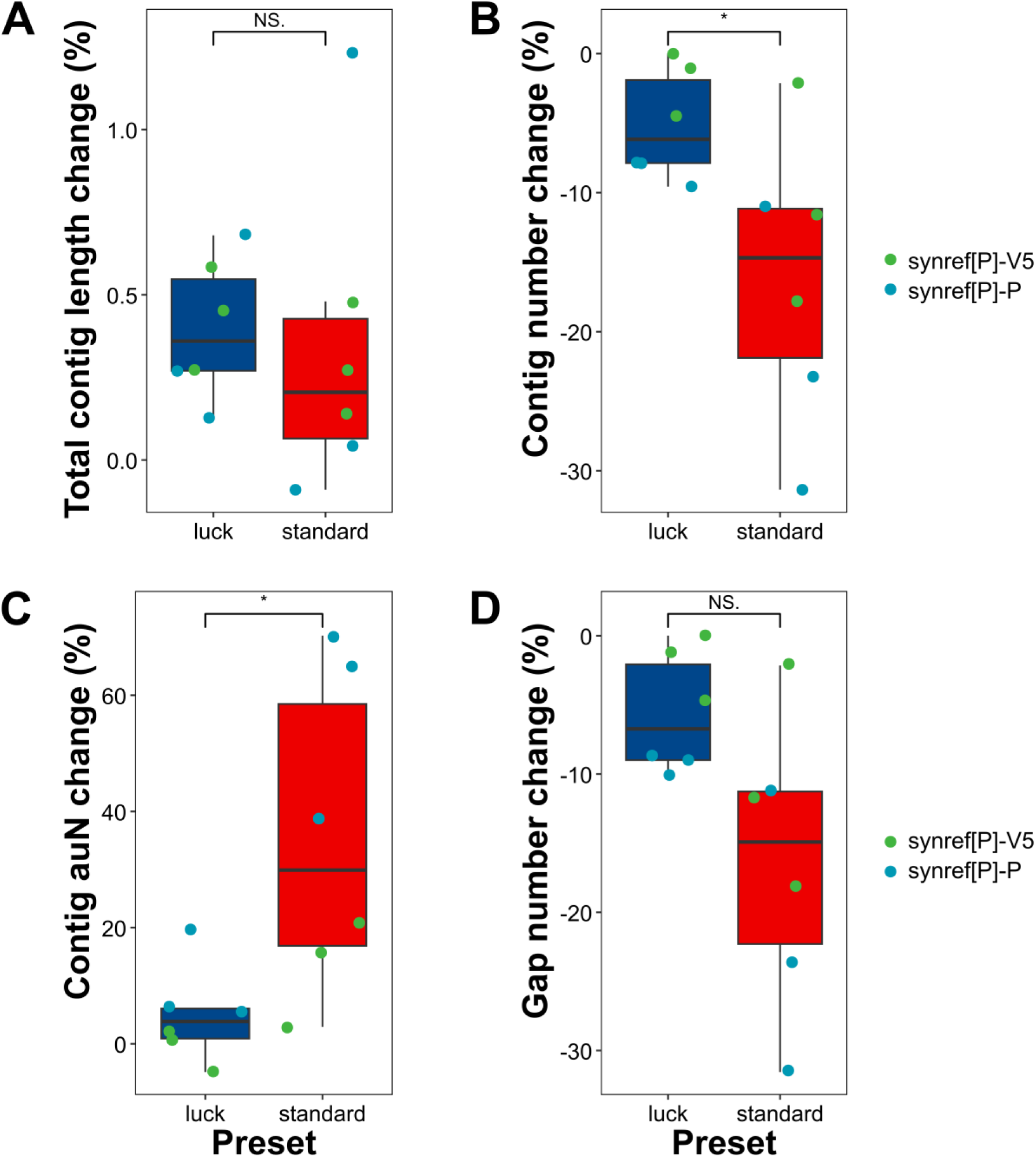
Assessment of the contig contiguity-preserving effect of synref[P]s during reference-guided error correction under the “luck” and “standard” presets, based on percentage changes in total contig length (**A**), contig number (**B**), contig auN (**C**), and gap number (**D**) between assemblies guided by synref[P]s and those guided by V5 or P references. The data points with different colors show reference contrasts in metrics.

When percentage changes in the four contig contiguity metrics were calculated between assemblies guided by synref[P]s and those guided by V5 or P references, the advantages of synref[P]s in reducing contig fragmentation during reference-guided error correction were most evident under the “standard” preset (**Figure 4**). Although assemblies scaffolded with synref[P]s generally showed higher contig auN values and lower contig numbers than those guided by other references under both the “luck” and “standard” presets, the effect of synref[P]s on preserving contig contiguity was significantly greater in the “standard” preset than in the “luck” preset (*p* < 0.05). The maximum observed increases in contig auN and reductions in contig number were 65.05% and 31.56%, respectively (**Figure 4B & C**). Under both presets, changes in total anchored contig length between assemblies guided by synref[P]s and those guided by other references were not significant, with values generally ranging from 0 to 0.5% (**Figure 4A**).

Statistical comparisons were performed using independent *t*-tests with α = 0.05.

**Figure 5** provides a more detailed presentation of the results shown in **Figure 4**. Under the “luck” correction preset, increases in contig auN for assemblies guided by synref[P]s relative to those guided by the V5 reference ranged from 0.121 to 0.384 Mb. Notably, for B108 under the “luck” preset, the synref[P]-guided assembly exhibited an approximately 0.43 Mb lower contig auN than the assembly guided by V5 (**Figure 5A**). In contrast, when the stricter “standard” correction preset was applied, no reductions in contig auN were observed for assemblies guided by synref[P]s relative to V5, and the maximum auN increase reached approximately 1.3 Mb (**Figure 5B**). A similar pattern was observed for contig number. Under the “luck” preset, the maximum reduction in contig number when comparing synref[P]- and V5-guided assemblies was eight contigs, whereas under the “standard” preset, this reduction increased to a maximum of 150 contigs (**Figure 5**). The contig contiguity-preserving effect of synref[P]s during correction relative to V5 was also evident in total anchored contig length, which was consistently higher in synref[P]-guided assemblies under both presets, with length increases ranging from 1.022 to 4.148 Mb (**Figure 5**).

**Figure 5:**
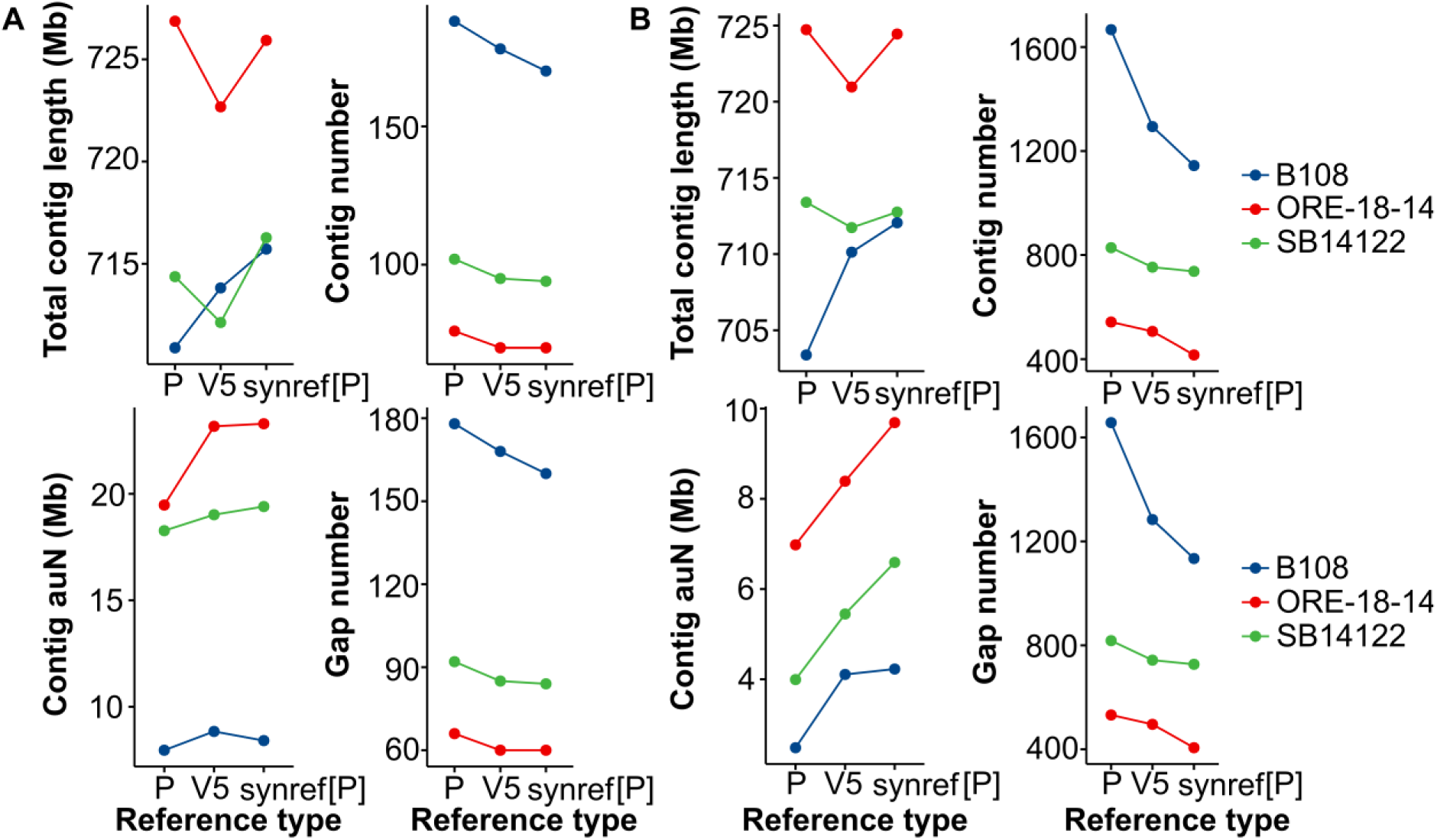
Comparison of in-chromosome contig contiguity metrics for assemblies scaffolded using synref[P], V5, and P references under the “luck” (A) and “standard” (B) presets.

The P references were shown in **Figure 5** to be the least effective reference type for target contig correction and scaffolding. Assemblies built using these references consistently exhibited lower contig auN values, higher contig numbers, and higher gap numbers than assemblies guided by synref[P]s or V5 under both the “luck” and “standard” presets. Despite being gapless genomes, the P references were not effective in reducing break number (i.e., the difference in total contig counts before and after correction) during contig correction prior to scaffolding (**Supplementary Figures S1** & **S2**).

With respect to structural correctness metrics, the examined assemblies showed only minor differences in QV, R-AQI, and S-AQI across most cases, with the exception of QV values under the “standard” preset. For the B108 accession, the QV of the assembly scaffolded using the P reference was approximately five units lower than those scaffolded using synref[P] and V5. In SB14122, the QV of the synref[P]-guided assembly decreased from 46.02 to 43.39 relative to the V5-guided assembly. However, this reduction in QV was modest, representing a 5.71% decrease compared with the V5 case (**Figure 6**).

**Figure 6:**
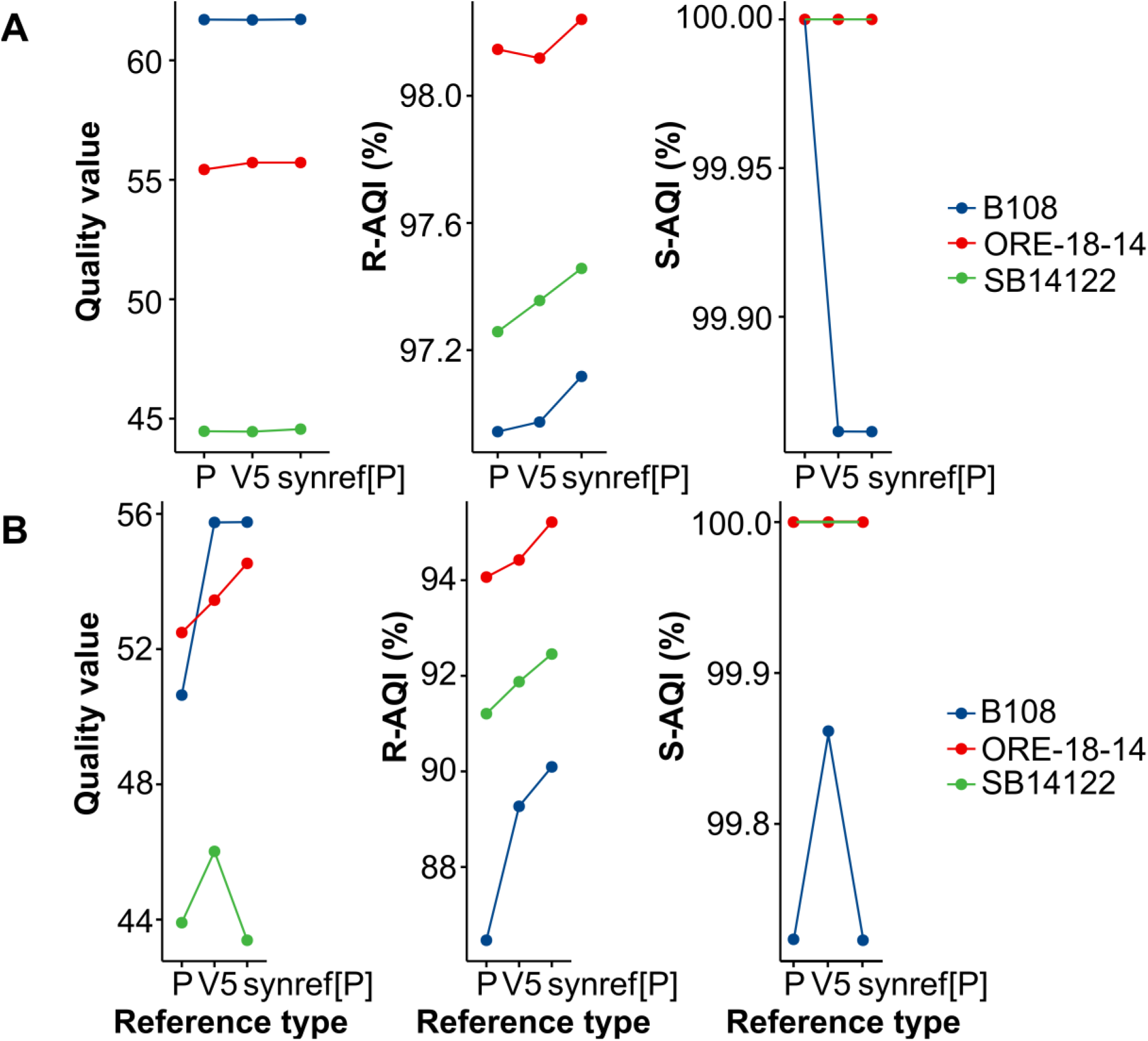
Comparison of structural correctness metrics for assemblies scaffolded using synref[P], V5, and P references under the “luck” (A) and “standard” (B) presets.

To further assess the structural correctness of all *Sbi* assemblies generated by noHiC, dot plots were constructed to illustrate syntenic relationships between V5 and the noHiC assemblies. Across all three accessions, the “luck” preset failed to correct large-scale misassemblies, manifested as false interchromosomal translocations. To find the reasons, we mapped reads to the scaffolded chromosomes and observed secondary alignments near the boundaries of these misjoins, which likely interfered with both read-clip-based and contig-alignment-based chimeric contig breaking processes. These misassemblies were caused by the contig assembler since sequences from different chromosomes were joined together in single chimeric contigs (**Supplementary Figures S3-S5**). These translocations occurred between chromosomes 4 and 1 in SB14122 (**Figure 7A**), chromosomes 5 and 6 and chromosomes 4 and 7 in B108 (**Figure 7B** & **C**), and chromosomes 4 and 7 in ORE-18-14 (**Figure 7D**). The misassemblies were corrected or markedly reduced when the “standard” preset was applied.

**Figure 7:**
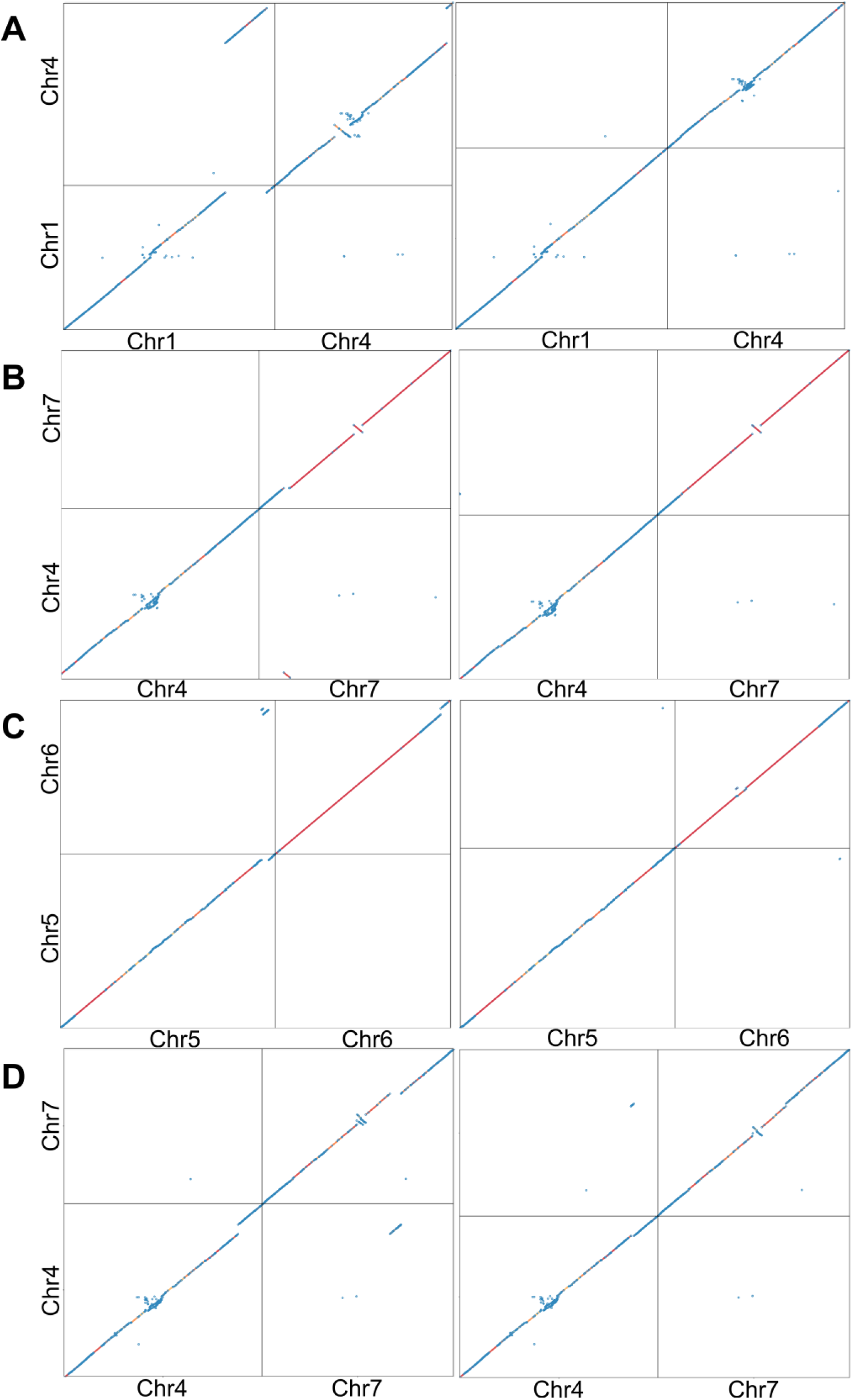
Examination of syntenic relationships between noHiC assemblies (y-axes) and V5 (x-axes), showing false interchromosomal translocations under the “luck” preset (left panels) for SB14122 (**A**), B108 (**B**, **C**), and ORE-18-14 (**D**). Misassemblies corrected by the “standard” preset are shown in the right panels. Dot plots are shown for assemblies guided by synref[P]s as representative examples.

### Synref’s Positive Effects in Contig Scaffoldings of Different Species

We scaffolded genomes from four species, including *A. thaliana* CAMA-C-2 (*Ath* CAMA-C-2),

*S. virgatum* (*Svi*), *G. max* cv. Wm82-ISU-01 (*Gma* cv. Wm82-ISU-01), and *H. vulgare* cv. Foma (*Hvu* cv. Foma) using noHiC with two reference types, namely a synref and a conventional reference genome (**Table 1**), to assess whether the advantages of synrefs could be observed across different plant species. The conventional references listed in **Table 1** are NCBI reference genomes or the subsequent versions of those references. The Jack genome was selected as a gapless and genetically close reference for the contig scaffolding of *Gma* cv. Wm82-ISU-01. Public original assemblies constructed using Hi-C data (or manual curation in the case of *Ath*) were selected as controls for evaluating the assemblies generated by noHiC. Comparing assemblies guided by synrefs, ordinary references, and controls will determine if selecting an assembly labeled as a “reference” in NCBI reliably guides a target genome scaffolding.

As shown in **Figures 4**, **5**, & **7**, although the “standard” preset effectively highlighted the advantages of synrefs for contig scaffolding and corrects contig misjoins that were not detected by relaxed presets such as “luck,” it also substantially reduced contig auN. This indicates that, unless hard-to-detect chimeric contigs are present or the reference genome has a highly similar sequence organization to the target genome, the “standard” preset should not be the initial choice, despite being the default option. Accordingly, in this computational evaluation of noHiC involving target assemblies from four plant species, the *nohic-asm* sub-script was executed using the “luck” preset with unpatched synrefs. Because most conventional references were not the genomes most closely related to the target assemblies, the “luck” preset was selected instead of the “standard” preset to avoid bias in favor of synrefs that could arise if assemblies guided by conventional references became excessively fragmented under strict correction.

Consistent with the phylogenetic relationships observed for the three sorghum assemblies, synrefs for the four examined plant species were also the closest references to their respective target genomes based on genetic distance (**Figure 8**). This advantage of synrefs was observed across pangenomes with varying numbers of haplotypes, ranging from 10 in barley to 48 in *Arabidopsis*.

**Figure 8:**
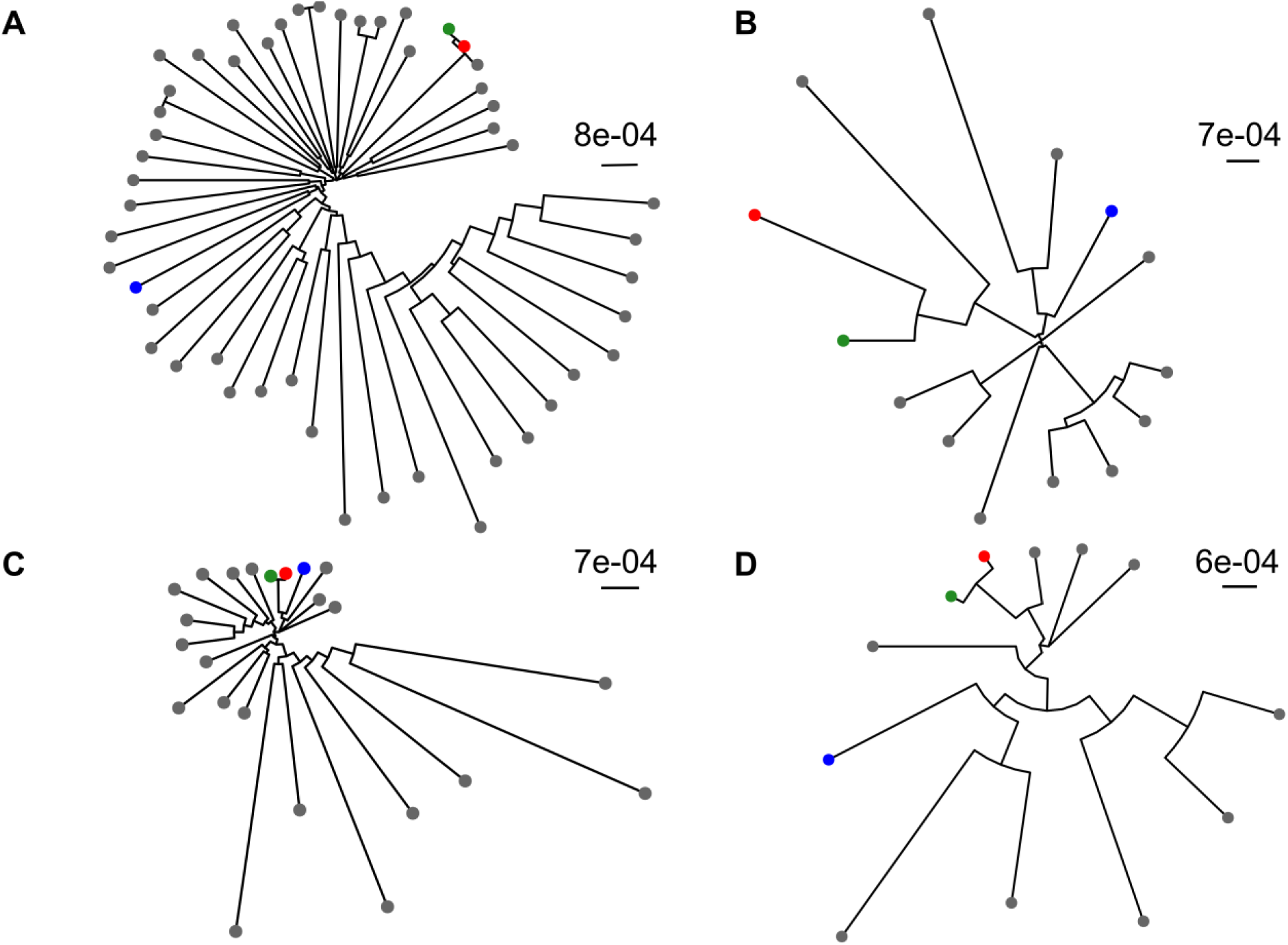
Visualization of NJ phylogenetic trees for *Ath* (**A**), *Svi* (**B**), *Gma* (**C**), and *Hvu* (**D**). Red, green, blue, and grey dots denote the target contig-level assemblies, synrefs, ordinary references, and other assemblies in the pangenome collections, respectively.

As shown in **Figure 9A and B**, scaffold contiguity metrics, including total scaffold length and scaffold auN, were largely comparable between assemblies generated using synrefs and those generated using conventional reference genomes.

**Figure 9:**
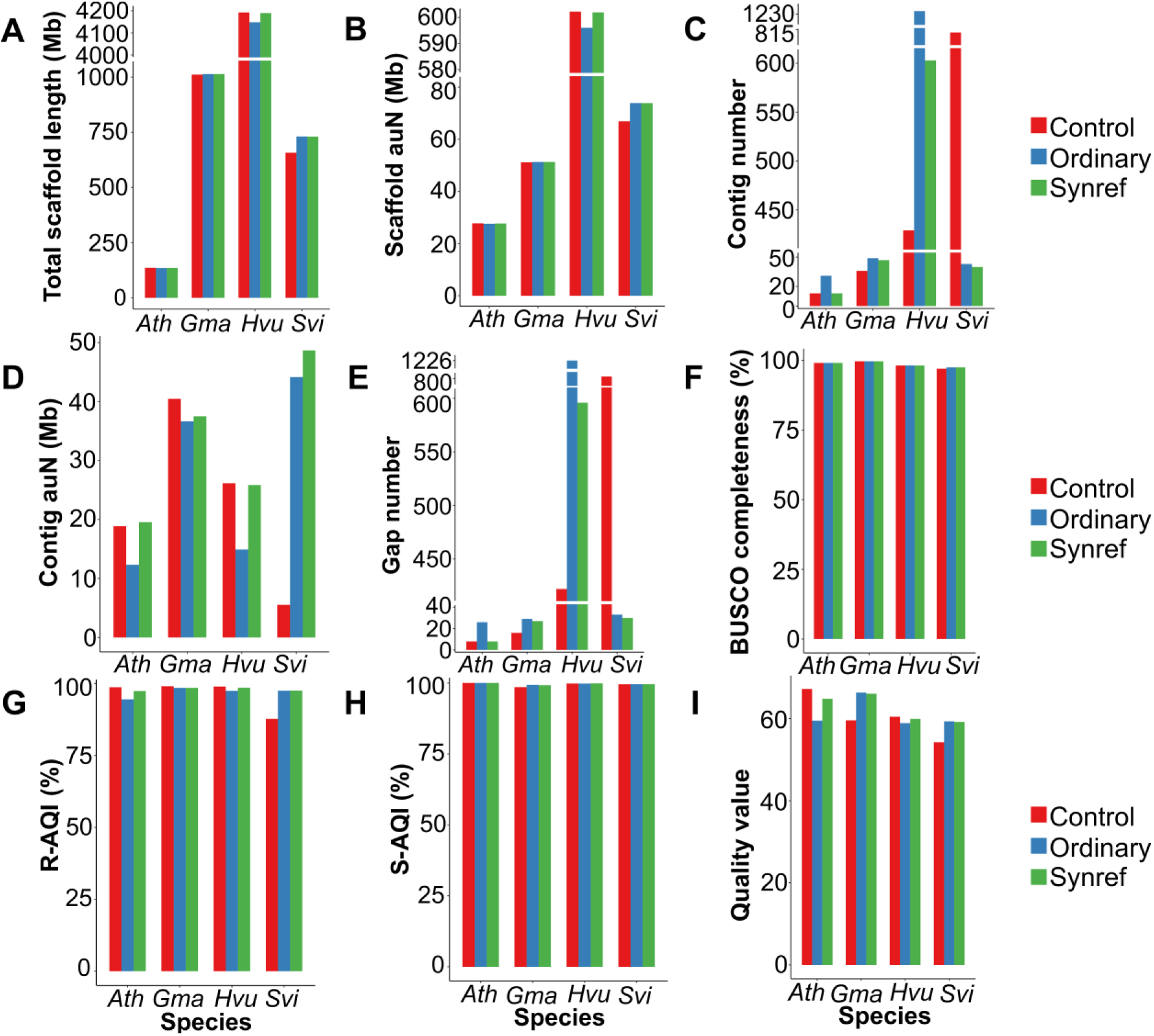
Comparison of in-chromosome contiguity and structural correctness metrics among conventionally guided assemblies (category “Ordinary”), synref-based assemblies (category “Synref”), and control assemblies (category “Control”) for four plant species (*Ath*, *Svi*, *Gma*, and *Hvu*).

Once again, the advantages of synrefs were most clearly reflected in in-chromosome contig contiguity metrics prior to gap closing. For contig auN, three of the four species (excluding *Gma*) showed clear increases ranging from 10.25% to 73.20% when synrefs were used instead of conventional references (**Figure 9D**). In the case of *Ath* CAMA-C-2, the synref-guided assembly exhibited contig and gap numbers comparable to those of the control assembly and 2.38-fold and 3.25-fold lower, respectively, than the assembly guided by TAIR10.1 (**Figure 9C** & **E**).

*Hvu* cv. Foma was the species in which synref showed the strongest contiguity-preserving effects. Specifically, noHiC assemblies were more fragmented than the control assembly (**Figure 9C, D** & **E**). Assemblies constructed using the Morex v3 reference genome had contig and gap numbers approximately three times higher than those of the control. In contrast, the synref maintained a strong contiguity-preserving effect, reducing contig and gap numbers by more than 51% relative to the Morex v3-guided assembly (**Figure 9C** & **E**).

*Svi* represented a distinct case in which noHiC assemblies showed higher contiguity than the Hi-C-based control (*S. virgatum* 1.0). The noHiC assembly guided by the conventional BTx623 v5 reference exhibited nearly an eightfold increase in contig auN compared with the control, along with 19-fold and 24-fold reductions in contig number and gap number (**Figure 9C, D** & **E**), respectively. The synref-guided *Svi* assembly showed even greater improvements, with nearly ninefold increases in contig auN relative to the public Hi-C-based control assembly (**Figure 9D**).

Consistent with observations from the three sorghum assemblies described above, under the “luck” preset, *Gma* was the only species for which synref did not show a clear advantage over a gapless reference genome that is genetically close to the Wm82-ISU-01 genome (genetic distance: 0.16%). In this case, only minimal differences were observed between synref- and conventional reference-guided assemblies, including a 2.40% increase in contig auN, a 4.08% reduction in contig number, and an approximately 7% reduction in gap number (**Figure 9C, D & E**).

In most cases, structural correctness metrics did not differ substantially between assemblies guided by synrefs and those guided by conventional references, with the exception of *Ath*, for which the QV of the synref-guided assembly was approximately 8.96% higher than that of the assembly guided by the conventional reference TAIR10.1. BUSCO completeness values for the noHiC assemblies were comparable to those of the control assemblies (**Figure 9F, G, H & I**).

With respect to synteny analyses between noHiC assemblies and their public counterparts, the synref-guided assembly of *Ath* CAMA-C-2 showed better agreement with the control than the assembly generated using the TAIR10.1 reference, which exhibited false chromosomal translocations between chromosomes 1 and 5. In the case of *Svi*, both noHiC assemblies contained longer centromeric regions than the *Svi* 1.0 assembly. *Gma* was the only species in which a sequence misarrangement was detected, occurring on chromosome 17 in the synref-guided assembly. This might be due to the *Gma* unpatched synref (see the scaffolding results with the patched one in **Figure 12**). For *Hvu*, both noHiC assemblies showed good syntenic correspondence with the public *Hvu* cv. Foma assembly (Figure 10**, Supplementary Figures S6-S9**).

**Figure 10:**
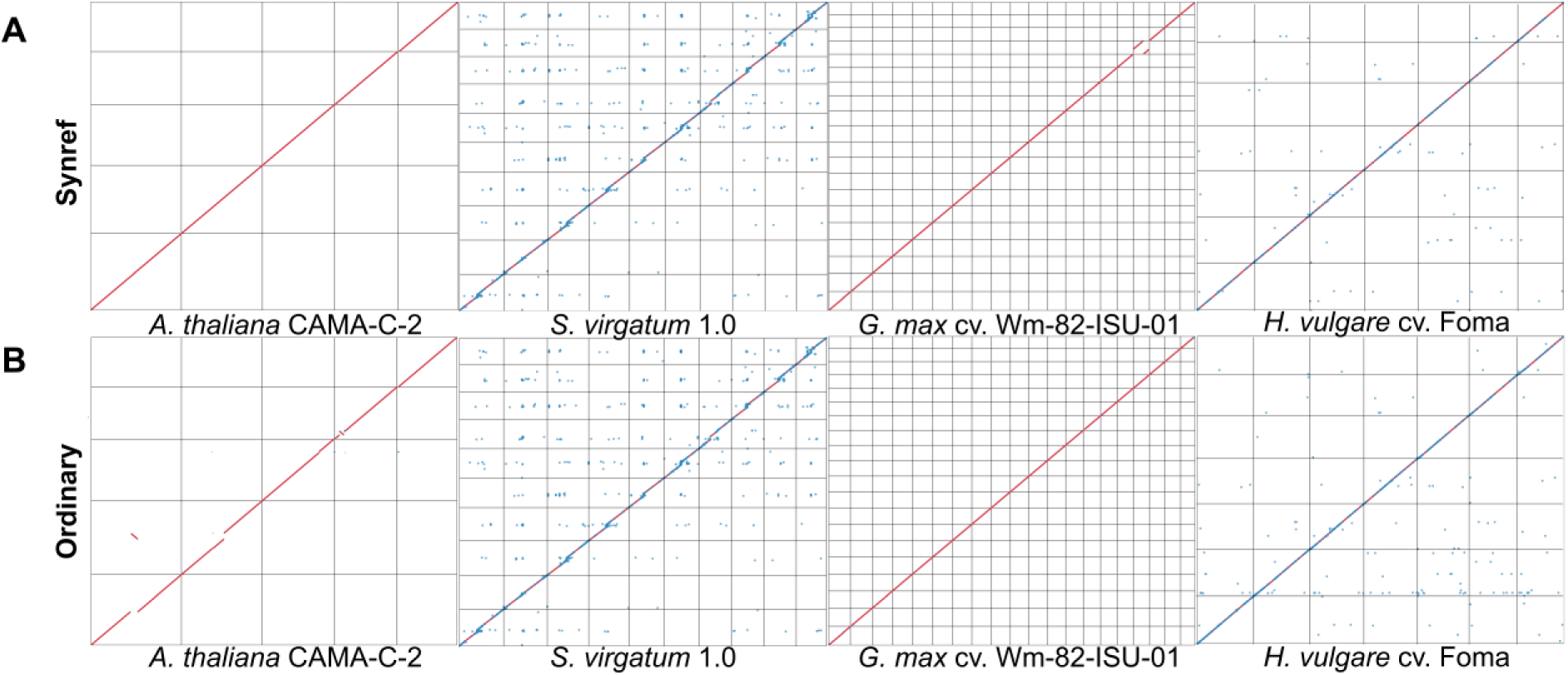
Visualization of dot plots showing whole-genome alignments between synref-based assemblies (category “Synref”) (**A**) or conventionally guided assemblies (category “Ordinary”) (**B**) and control genomes (x-axes) for four plant species. Each square containing a diagonal represents a chromosome. Chromosomes are ordered according to their lengths in the control assemblies.

### The Flexibility of noHiC Pipeline when Combined with a Fast Scaffolder

Recognizing that each contig correction step in *nohic-asm* requires substantial computational time, we replaced *nohic-asm* with ntJoin, an ultra-fast scaffolder, to correct and scaffold contigs from *Ath* CAMA-C-2, *Svi*, *Gma* cv. Wm82-ISU-01, and *Hvu* cv. Foma (**Table 2**). This analysis aimed to assess whether *nohic-refpick* and *nohic-clean* can be flexibly combined with other fast scaffolding algorithms to improve assembly quality compared with the use of a single reference. Because ntJoin does not rely on sequence alignment and breaks contigs without read-based validation, the scaffolding of *Svi*, *Gma* cv. Wm82-ISU-01, and *Hvu* cv. Foma were guided by patched synrefs (synref[P]s) (**Table 1**) to ensure that gaps did not limit the benefits of synrefs. The synref for *Ath* CAMA-C-2 was left unpatched, as it exhibited higher contiguity than TAIR10.1.

**Table 2.** Computational resources required for contig correction and scaffolding using *nohic-asm* and ntJoins.

| Scaffolding tool | Species | Used CPU core number | Memory use (GB) | Run time (d-hh:mm:ss) |
| --- | --- | --- | --- | --- |
| *nohic-asm* | *Ath* | 21 | 68.28 | 01:00:49 |
| *nohic-asm* | *Svi* | 40 | 207 | 10:39:08 |
| *nohic-asm* | *Gma* | 36 | 193.12 | 09:58:34 |
| *nohic-asm* | *Hvu* | 35 | 259.32 | 2-23:40:20 |
| ntJoin | *Ath* | 1 | 1.76 | 00:01:45 |
| ntJoin | *Svi* | 2 | 12.29 | 00:04:52 |
| ntJoin | *Gma* | 3 | 7.15 | 00:02:39 |
| ntJoin | *Hvu* | 3 | 17.40 | 00:11:19 |

**Note:** Because assemblies were performed twice for each species using a conventional reference and a synref or synref[P], the values reported correspond to the assemblies with higher computational demands.

As shown in **Supplementary Table S2**, ntJoin assemblies generally exhibited higher contig counts than those generated by noHiC, indicating that ntJoin introduced a greater number of contig breaks. However, in contrast to assemblies scaffolded using *nohic-asm*, ntJoin assemblies guided by a synref or synref[P] (synref/synref[P]) exhibited more pronounced improvements compared to conventionally guided assemblies in scaffold contiguity metrics, including total scaffold length (Mb) and scaffold auN (Mb). Specifically, when synref/synref[P]-guided assemblies were compared with those based on conventional references, percentage increases in total scaffold length and scaffold auN ranged from 3.52% to 39.34% and from 2.33% to 16.20%, respectively (**Figure 11A & B**).

Despite being applied with a different scaffolding algorithm, synrefs/synref[P]s continued to show strong effects in limiting contig fragmentation during correction. Specifically, synref/synref[P]-guided assemblies for all four species exhibited higher contig auN values than those guided by conventional references. For all species, the increases in contig auN were substantial, ranging from 39.55% to 57.77% (**Figure 11D**). Consistent with the contig auN results, the use of synrefs (or synref[P]s) in ntJoin-based scaffolding also reduced contig and gap numbers for *Ath*, *Gma*, and *Hvu*, with reductions of 76.22%, 36.61%, and 51.92% in contig number and 82.26%, 54.05%, and 53.27% in gap number, respectively (**Figure 11 C** & **E**). Notably, for *Gma* cv. Wm82-ISU-01, although both the synref[P] and the conventional reference (the Jack genome) were gapless, assemblies derived from the synref[P] still showed higher contig auN values, as well as lower contig and gap numbers, than assemblies guided by the genetically close, gapless Jack reference (**Figure 11C, D** & **E**).

BUSCO completeness values, which have been reported to show little sensitivity to reference genome choice in previous comparisons, were higher in ntJoin assemblies guided by synrefs/synref[P]s. Specifically, BUSCO completeness values for synref/synref[P]- and conventional-reference-guided assemblies were 99.1% and 93.9% for *Ath* CAMA-C-2, 77.5% and 60% for *Svi*, 96.8% and 92.5% for *Hvu* cv. Foma, and 99.6% and 98.8% for *Gma* cv.

Wm82-ISU-01, respectively (**Figure 11F**).

Regarding structural correctness metrics, ntJoin assemblies derived from synrefs/synref[P]s generally exhibited higher R-AQI, S-AQI, and QV values than those guided by conventional references (**Figure 11G, H** & **I**). The most pronounced improvements in structural correctness for synref/synref[P]-guided assemblies were observed for the QV metric, with increases of 25.03, 6.22, 5.58, and 9.86 units for *Ath*, *Svi*, *Hvu*, and *Gma*, respectively (**Figure 11**).

**Figure 11:**
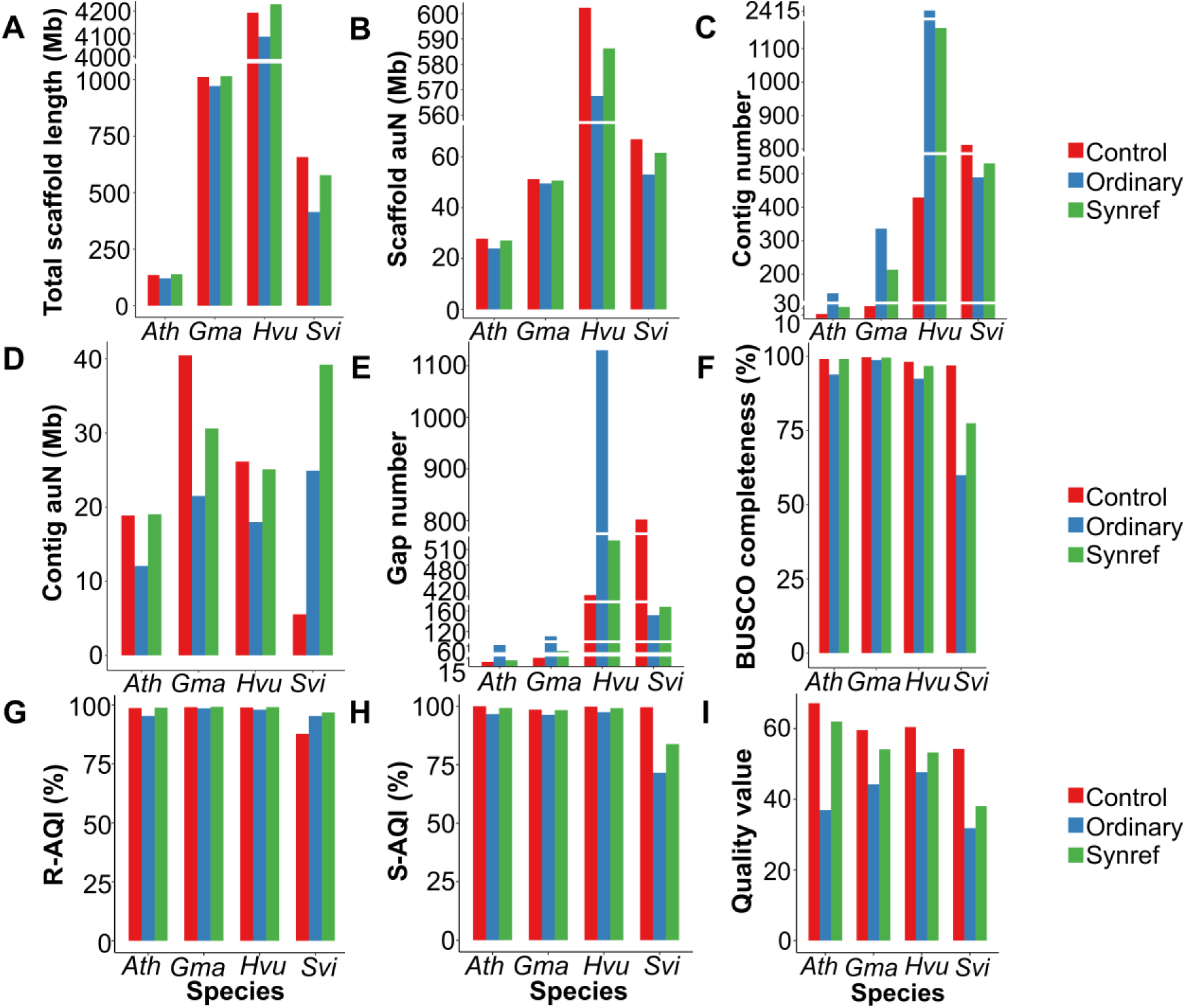
Comparison of contiguity and structural correctness metrics (calculated from whole assemblies) among ntJoin assemblies constructed with different references for four plant species (*Ath*, *Svi*, *Gma*, and *Hvu*). The categories “Ordinary”, “Synref”, and “Control” are for conventionally guided assemblies, synref/synref[P]-based assemblies, and original public assemblies, respectively.

ntJoin assemblies derived from synrefs (or synref[P]s) showed better syntenic agreement with the control assemblies than those guided by conventional references in three of the four species examined (*Ath* CAMA-C-2, *Gma* cv. Wm82-ISU-01, and *Hvu* cv. Foma) (**Figure 12A**). Specifically, in the *Ath* case, the assembly guided by the TAIR10.1 reference showed a shortened chromosome 2 and false chromosomal translocations between chromosomes 1 and 4. In *Hvu* cv. Foma, the assembly scaffolded using the Morex v3 reference exhibited false chromosomal translocations involving chromosomes 2H, 3H, 4H, 6H, and 7H. For *Gma* cv.

**Figure 12:**
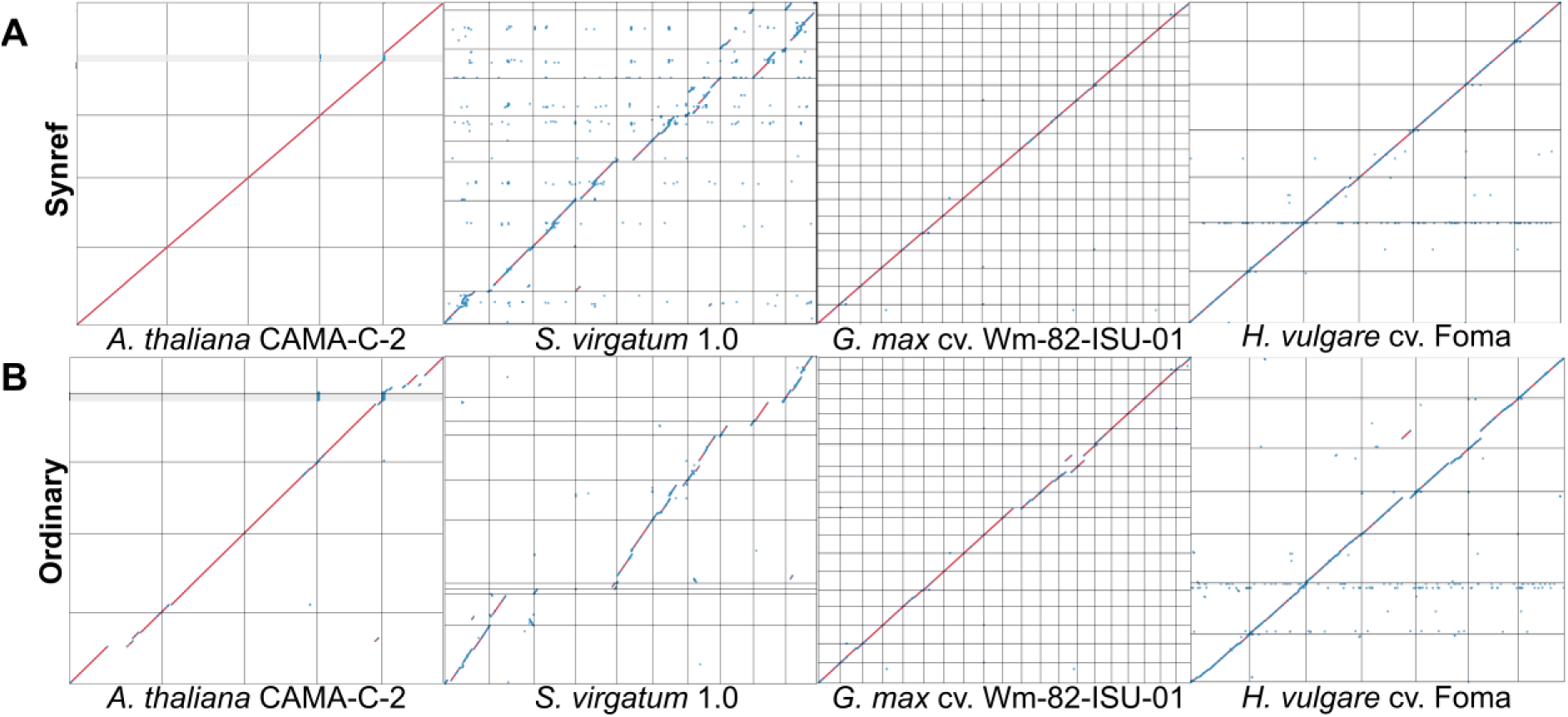
Visualization of dot plots showing whole-genome alignments between synref/synref[P]-based assemblies (category “Synref”) (**A**) or conventionally guided ntJoin assemblies (category “Ordinary”) (**B**) and control genomes (x-axes) for four plant species. Each square containing a diagonal represents a chromosome. Chromosomes are ordered according to their lengths in the control assemblies.

Wm82-ISU-01, the assembly derived from the Jack reference showed a shortened chromosome 9 and a large false chromosomal translocation between chromosomes 3 and 7 (**Figure 12B**). *Svi* was the only case in which assemblies guided by both the synref[P] and the BTx623 v5 reference did not align well with the control public assembly (*Svi* 1.0). However, the diagonal pattern, showing sequence agreement between ntJoin assemblies and the controls, is more obvious in the synref[P]-based assembly compared to the ordinary-guided one. (**Figure 12, Supplementary Figures S10-S13**).

## DISCUSSION

In this study, we designed and evaluated the noHiC pipeline, which was developed to apply multiple strategies to ensure the quality of reference-guided contig scaffolding. The pipeline removes contaminant DNA sequences originating from non-target taxonomic groups present in the input contigs, thereby reducing the risk of false interpretations in evolutionary and interspecific lateral gene transfer analyses based on the scaffolded genomes (80–82). In addition, because noHiC does not rely on optical mapping (83) or Hi-C sequencing data (84), which are commonly used for validation and correction of contig misjoins (43, 85), three complementary contig polishing approaches were implemented in *nohic-asm* to correct small errors and to break potentially misassembled junctions (18, 43, 44). However, because contig misjoin correction relies on alignment to a reference genome, noHiC may incorrectly break truly contiguous contigs. To address this limitation, we created *nohic-refpick* to generate a personalized reference (synref) for each target assembly using a pangenome graph-based haplotype sampling algorithm (30). The synref is an artificial genome composed of 10-kb fragments that best match the target genome, enabling the incorporation of genetic information from large numbers of reference genomes in a pangenome graph for scaffolding - up to 48 genomes in the case of *A. thaliana* - without directly aligning to each reference. The positive effects of synrefs in alleviating reference bias primarily involve preservation of contig contiguity during reference-guided correction, which are consistently demonstrated in the analyses described below.

We evaluated the use of the noHiC pipeline with different synrefs derived from an 11-haplotype pangenome graph for scaffolding three sorghum accessions, namely SB14122, B108, and ORE-18-14. Analysis of the relationships among each target accession, the synref, the V5 reference, the P reference, and other accessions included in the pangenome graph showed that a single pangenome graph can be reused to generate genetically best-fitting references for multiple assemblies of the same species. Using two contig correction presets (“luck” and “standard”), we found that the contig break-prevention effect of synrefs was most evident when contigs from a target genome were constrained to adopt sequence arrangements similar to those of the reference genome, as in the “standard” preset.

Assemblies generated using a strict correction preset can exhibit lower contig contiguity than those produced under more relaxed presets. However, strict correction is required in cases where secondary read alignments bypass relaxed correction strategies, leaving misjunctions in contigs undetected. Under such conditions, the contig contiguity-preserving effects of synrefs are particularly advantageous. Comparisons of synrefs with other reference types, including the genetically closest reference to the target genomes aside from synrefs and gapless sorghum genomes, under the strict contig correction preset showed consistent improvements in contig contiguity metrics when synrefs were used for target genome scaffolding. Therefore, relative to the use of synrefs, relying solely on a closely related or gapless genome as a reference is insufficient to overcome reference bias during contig correction and scaffolding.

The demonstrated reusability of pangenome graphs for generating best-fit references distinguishes noHiC from other multi-reference scaffolders. Users can construct a high-quality pangenome graph that captures extensive genetic diversity across multiple available genomes and subsequently use this graph to generate different synrefs for future scaffolding projects. By employing *nohic-refpick* for synref generation, users are not required to perform reference weight optimization, as is necessary in ntJoin (26) (https://github.com/bcgsc/ntJoin). In addition, unlike Ragout2-based scaffolding, noHiC does not require repeated updating of alignments between the target genome and multiple references whenever a new assembly project is initiated. The use of *nohic-refpick* reduces the alignment task from multiple genome-to-genome alignments to a single alignment between the synref and the target genome (20, 27) (https://github.com/ComparativeGenomicsToolkit/cactus/blob/master/doc/updating-alignments.md). Similarly, in contrast to multi-CSAR (25), noHiC eliminates the need to repeatedly align each new target genome against multiple individual references, as the alignment is performed only against the generated synref.

The prevention of contig breakage through the use of synrefs during reference-guided correction and scaffolding was also clearly observed in most of the plant species tested (*Ath*, *Svi*, and *Hvu*), which include both monocots and dicots and span a wide range of genome sizes (approximately 135 Mb to 4.2 Gb), when scaffolding results based on synrefs were compared with those based on NCBI references (or the subsequent NCBI reference version). These results indicate that the noHiC pipeline has the potential to be applied broadly to contig scaffolding across diverse plant species. In addition, the findings suggest that straightforward use of an NCBI reference alone as a guide for contig correction and scaffolding is insufficient to ensure assembly contiguity during contig correction. However, under relaxed contig correction conditions, where correction relies on sequence alignments with an aligner that does not require long exact seed matches between target and reference genomes (e.g., minimap2), synrefs may not outperform a highly contiguous conventional reference that is genetically close to the target genome, as observed in three sorghum accessions and *Gma* cv. Wm82-ISU-01.

In terms of structural correctness, noHiC has the capacity to generate reference-guided assemblies that are structurally consistent with public Hi-C-based or manually curated genomes (controls). Assemblies that are highly syntenic to public controls can be obtained using unpatched synrefs (in most cases) or a gapless reference closely related to the target genome. These observations are reproducible and remain evident when patched synrefs are applied with an alternative scaffolding tool. Thus, noHiC provides a reliable approach for generating chromosome-scale assemblies while reducing the need for Hi-C sequencing data.

The running time of *nohic-asm* is substantially increased by four steps that require read mapping to contigs or scaffolds, including CRAQ, Inspector, *ragtag.py correct*, and TGSGapcloser. As a result, users may prefer to employ an ultra-fast scaffolder such as ntJoin for contig correction and scaffolding when computational time is a limiting factor. In such cases, noHiC remains applicable, as users can leverage publicly available pangenome graphs and apply *nohic-refpick* to generate a best-fit reference for contig correction and scaffolding. Our analyses showed that combining ntJoin and noHiC’s subscripts (*nohic-clean* and *nohic-refpick*) markedly reduced runtime and computational resource requirements for these tasks. However, ntJoin assemblies generally exhibited higher contig counts (a sign of increased contig breakage during reference-guided correction) than those generated by noHiC. This indicates that, although the read-mapping-based contig-breaking steps in noHiC substantially increase computational time and resource requirements, they play an important role in limiting excessive contig fragmentation. Consequently, users should carefully consider the trade-off between scaffolding time and assembly quality before combining noHiC sub-scripts with alternative scaffolding tools.

In addition, assembly results obtained by combining *nohic-refpick* with the non-alignment-based correction and scaffolding strategy of ntJoin indicate that the advantages of synrefs are not restricted to the noHiC pipeline. Specifically, ntJoin assemblies guided by synrefs showed clear improvements over those guided by conventional references across multiple metrics and syntenic analyses in all of the plant species tested. These results demonstrate the flexibility and effectiveness of *nohic-refpick* when used in combination with alternative scaffolding methods. Notably, in the case of *Gma* cv. Wm82-ISU-01, the synref-guided assembly outperformed the Jack reference, despite the latter being a gapless genome genetically close to the target. Thus, a gapless genome closely related to the target is not necessarily the optimal reference for all scaffolding algorithms. *Svi* was the only case in which new assemblies were of lower quality than the control assembly (*Svi* 1.0), indicating that ntJoin is highly species-specific and may not be suitable for scaffolding even when the reference genome originates from a closely related but distinct species.

Despite the advantages described above, synref-guided contig scaffolding has a limitation. The haplotype sampling algorithm selects the best-fitting 10-kb DNA segments from haplotypes within a pangenome graph and combines them to create a synref for the target genome (30), but it does not include a mechanism to verify the correctness of these segments. As a result, synrefs may inherit assembly errors present in haplotypes derived from low-quality genomes, potentially affecting the outcome of reference-guided scaffolding.

Therefore, users should ensure that the individual reference genomes incorporated into a pangenome graph are of high quality before applying noHiC for contig scaffolding. For example, running *nohic-eval* to assess the quality of public genomes included in the pangenome graph is an effective approach to verify their suitability.

## ACKNOWLEDGEMENTS

The graphical abstract was created by Biorender (https://BioRender.com/2wwli91).

## AUTHOR CONTRIBUTIONS

An Nguyen-Hoang: Conceptualization, Formal analysis, Methodology, Validation, and Writing-original draft. Kübra Arslan: Methodology and Writing-review & editing. Venkataramana Kopalli: Formal analysis and Writing-review & editing. Steffen Windpassinger: Resources. Dragan Perovic: Resources and Writing-review & editing. Andreas Stahl: Resources. Agnieszka Golicz: Conceptualization, Validation, Supervision, and Writing-review & editing.

All authors have read and approved the final manuscript.

## SUPPLEMENTARY DATA

### Supplementary Figures

Break counts introduced by noHiC during reference-guided contig correction using the “luck” (**Figure S1**) and “standard” (**Figure S2**) presets for three sorghum accessions with three reference types (V5, P, and synref[P]). Read mapping at the boundaries of false interchromosomal translocations in the SB14122 (**Figure S3**), B108 (**Figure S4**), and ORE-18-14 (**Figure S5**) noHiC sorghum assemblies (“luck” correction preset). Whole-genome alignments between conventionally guided or synref-guided noHiC assemblies and the corresponding public control assemblies for *Ath* (**Figure S6**), *Svi* (**Figure S7**), *Gma* (**Figure S8**), and *Hvu* (**Figure S9**). Whole-genome alignments between conventionally guided or synref-guided ntJoin assemblies and the corresponding public control assemblies for *Ath* (**Figure S10**), *Svi* (**Figure S11**), *Gma* (**Figure S12**), and *Hvu* (**Figure S13**).

### Supplementary Tables

Genome assemblies included in the pangenome graph collections and control genomes used in assembly tests for *Ath*, *Sbi*, *Svi, Gma*, and *Hvu* (**Table S1**). Comparison of contig counts between noHiC- and ntJoin-based whole assemblies, including pseudochromosomes and unplaced contigs (**Table S2**).

### Supplementary Data statement

Supplementary Data are available at NAR online.

## DATA AVAILABILITY

The pangenome graphs, contig assemblies, synrefs, scaffolded assemblies, and scripts underlying this study are available at Zenodo under DOI: 10.5281/zenodo.18720982.

